# *Eggerthella lenta* evades bacteriophage through reversible megabase-scale inversions of capsular polysaccharide gene clusters

**DOI:** 10.64898/2026.04.24.720693

**Authors:** Shenwei Zhang, Colin Buttimer, Kai R. Trepka, Kathy N. Lam, Luis A. Ramirez Hernandez, Paola Soto-Perez, Cecilia Noecker, Paolo Canigiula, Edwin F. Ortega, Jenny Lee, Lorenzo Ramirez, Gina Partipilo, Hugh B. Lawrence, Francesca Bottacini, Lorraine A. Draper, R. Paul Ross, Aidan Coffey, Andrey Shkoporov, Colin Hill, Peter J. Turnbaugh

**Author notes:** Correspondence to: Peter J. Turnbaugh,. Co-first authors (Shenwei Zhang, Colin Buttimer).

## Abstract

Bacteriophages are a promising tool for microbiome editing, yet their development has been constrained by limited insights into bacteriophage-host interactions within their shared mammalian body habitat. We isolated a lytic phage ΦKL11 that efficiently targets a disease-associated member of the human gut microbiota, *Eggerthella lenta*, during *in vitro* growth. However, ΦKL11 selects for a pre-existing and reversible bacteriophage-resistant sub-population in mice. Long-read sequencing revealed a massive genomic inversion event, representing >50% of the *E. lenta* genome, enriched in response to bacteriophage infection. Transcriptomics linked this inversion to the altered expression of three capsular polysaccharide synthesis (CPS) gene clusters and transmission electron microscopy confirmed differential capsule production. Finally, we show that ΦKL11 has a broad host range attributable to CPS and other strain-variable genes. These findings suggest a previously unrecognized strategy for phage evasion in the gut, involving megabase-scale genomic inversions and reversible capsule variation driving phage resistance.

## INTRODUCTION

The trillions of microorganisms found in and on the human body (the microbiota) and their aggregate genomes and metabolic activities (the microbiome) have been broadly implicated in the risk, progression, and treatment outcomes of numerous diseases^1–7^. However, definitive evidence for the necessity of individual gut microbial genes for specific host phenotypes remains out of reach due to the lack of precise and robust methods for removing individual species or genes from complex microbial communities. Instead, causality is typically established by experiments testing sufficiency, in which individual microbial genes are tested either alone or in defined microbial communities assembled in culture^8–11^.

Microbiome editing provides a potential solution^12,13^, leveraging the target specificity of CRISPR-Cas systems to precisely edit microbial communities^14,15^. The main challenge is to identify robust methods for *in situ* DNA delivery, prompting a reconsideration of bacteriophages (phages), which have evolved to do exactly that. Multiple *proof-of-concept* studies support the utility of bacteriophages for editing the *Escherichia coli* genome within the gastrointestinal tract^16–18^, but these tools have yet to be broadly implemented for other members of the human gut microbiota. In contrast to the rich history of phage biology in *E. coli*, we lack experimental data for the bacteriophages that infect the vast majority of prevalent human gut bacterial species. Tremendous progress has been made in mapping the human virome through sequencing-based methods, including sophisticated tools for host prediction^19,20^. Yet, phage isolation and experimental studies have lagged behind, especially with respect to studies in mammalian model organisms. Consistent with this gap, studies in defined and synthetic gut microbiota indicate that phage can persist and shape microbial function, but how individual gut commensals respond to sustained phage pressure *in vivo* remains unclear^21^.

We sought to address this knowledge gap by isolating bacteriophages that target the prevalent gut Actinomycetota *Eggerthella lenta*^22^. Recent work has emphasized the broad relevance of *E. lenta* for the metabolism of drugs^23,24^, plant polyphenols^25–27^, and host metabolites^28^. Furthermore, *E. lenta* has long been implicated in bacteremia^29^ and more recently, in inflammatory bowel disease^30–32^. Genome editing tools are also now available in *E. lenta*^33^, leveraging its endogenous Type I-C CRISPR-Cas system^19^. Thus, the discovery of bacteriophages that broadly target *E. lenta* promises to accelerate our ability to study the numerous ways in which *E. lenta* shapes disease risk and treatment outcomes.

Herein, we report the isolation and in-depth characterization of a novel broad host range lytic bacteriophage (ΦKL11) that infects *E. lenta.* Paired studies in cell culture and mouse models revealed a remarkable and unexpected mechanism through which *E. lenta* escapes phage targeting, in which a Mbp-sized reversible genomic inversion enables phase variable shifts in CPS gene expression that re-shape the cell surface. This system for genomic inversions has clear biotechnological implications, while also highlighting key challenges and opportunities for phage-based microbiome editing.

## RESULTS

### *E. lenta* evades bacteriophage in the gastrointestinal tract

We screened San Francisco wastewater samples collected in two consecutive years for bacteriophages capable of forming plaques on *E. lenta* DSM2243. Purification and genome sequencing identified 9 highly homologous (>95% nucleotide identity to a representative genome) tailed double-stranded DNA bacteriophages belonging to the class *Caudoviricetes,* which were consistently isolated across both years (**Table S1**). These phages also exhibited comparable genome sizes (range: 44,091 to 44,981 bp). Based on International Committee on Taxonomy of Viruses (ICTV) guidelines^34^, we assigned all 9 isolates to a single genus (tentative name: *Calivirus*), represented by ΦKL11, which served as the focus of all subsequent analyses. Annotation of the ΦKL11 genome identified 59 predicted ORFs, 46 of which belonged to protein clusters conserved across all nine phages at 90% amino acid identity, including 9 replication genes, 14 structural proteins, and 2 lysins (**Table S2**). ΦKL11 efficiently lysed *E. lenta* on solid (**Fig. 1A**) and in liquid (**Fig. 1B**) media, reaching titers of 9.6±0.05 log_10_ plaque-forming units (PFU) per mL. Transmission electron microscopy revealed that ΦKL11 exhibits the expected morphology of a tailed siphovirus (**Fig. 1C**).

**Figure 1.**
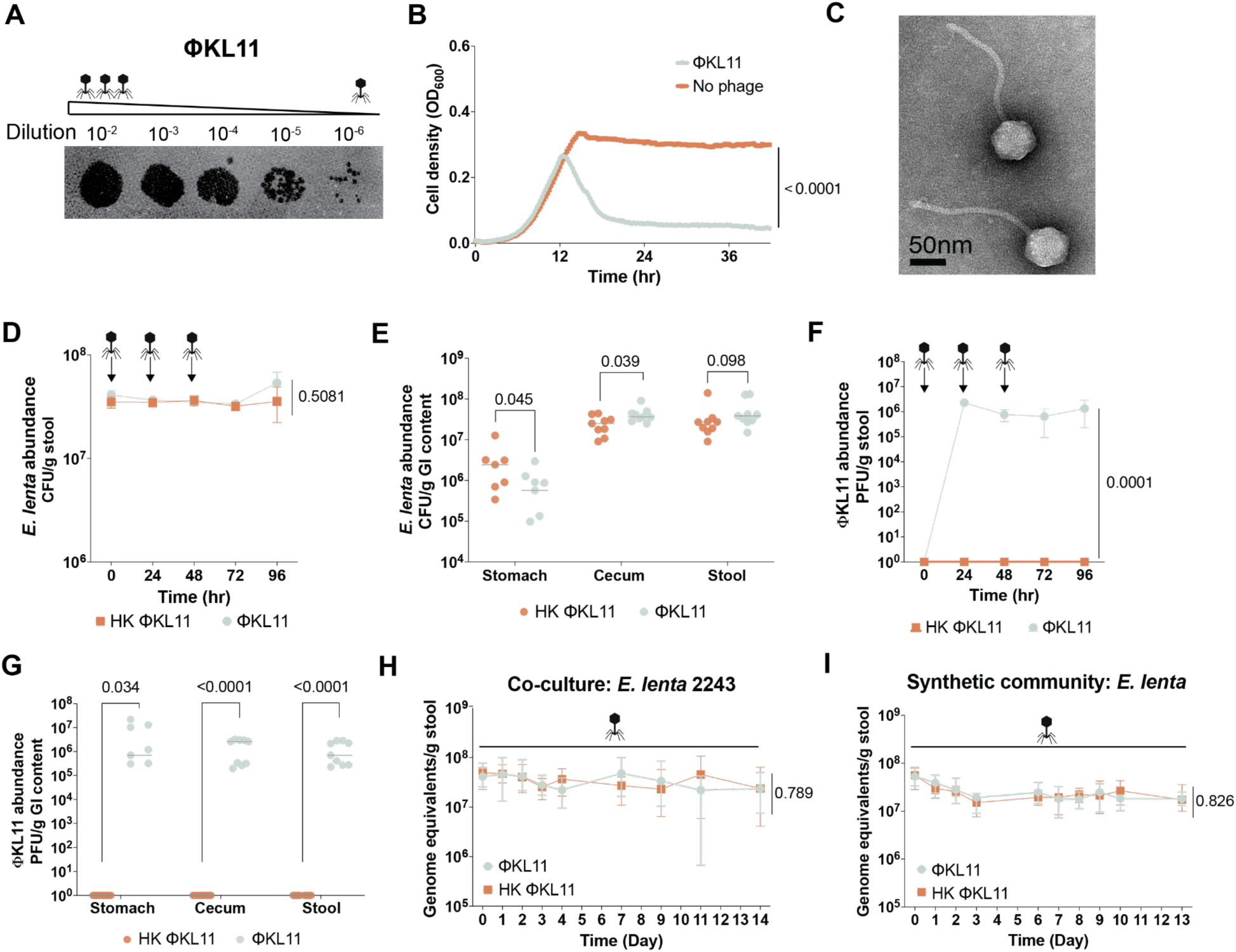
*E. lenta* evades ΦKL11 predation in the mouse gut. **(A–B)** ΦKL11 robustly lyses *E. lenta* DSM2243 on an agarose overlay **(A)** and in liquid BHI^CHAV^ medium **(B;** n=3 biological replicates). **(C)** Transmission electron micrograph of ΦKL11. Scale bar, 50 nm. **(D–G)** GF mixed-sex C57BL/6J mice were monoassociated with *E. lenta* DSM2243 for one week prior to phage treatment. Mice then received either heat-killed (HK) or active ΦKL11 by oral gavage for three consecutive days (n=9 mice/group). **(D,E)** *E. lenta* levels (CFU/g) and **(F,G)** ΦKL11 levels (PFU/g) in longitudinal stool samples **(D,F)** and endpoint contents from the stomach, cecum, and stool **(E,G). (H,I)** GF male BALB/c mice were colonized with defined bacterial communities and given either heat-killed (HK) or active ΦKL11 in drinking water for two weeks. **(H)** *E. lenta* DSM2243 quantification (qPCR) during co-colonization with the ΦKL11-resistant *E. lenta* DSM15644 (1:1 inoculation; n=9–10 mice/group). **(I)** *E. lenta* DSM2243 quantification (qPCR) during co-colonization with representative strains from three phyla: *Bacteroides thetaiotaomicron*, *Escherichia coli*, and *Clostridium innocuum* (1:1:1:1 initial inoculum; n=5 mice/group). Values are **(B)** mean, **(D,F,H,I)** mean±sem, and **(E,G)** median. *p*-values, two-way ANOVA **(B,D,F,H,I)** and Mann-Whitney U test **(E,G)**.

Despite this robust lytic activity *in vitro*, ΦKL11 had no detectable impact on *E. lenta* colonization *in vivo*. Germ-free (GF), mixed sex adult C57BL/6J mice were monocolonized with the ΦKL11-sensitive (ΦKL11^S^) *E. lenta* type strain DSM2243 and subsequently administered ΦKL11 or a heat-killed (HK) control by oral gavage once daily for three consecutive days (**Table S3A**). *E. lenta* abundance in stool remained stable for 4 days following phage administration (**Fig. 1D**) and was similarly maintained in endpoint stomach and cecal contents (**Fig. 1E**). In parallel, high titers of ΦKL11 were consistently detected throughout the experiment in stool samples (6.48 x 10^5^ - 2.28 x 10^6^ PFU/g stool; **Fig. 1F**) as well as in endpoint stomach and cecal contents (**Fig. 1G**), despite no phage dosing during the final 48 h. These findings were independently replicated in a separate experiment tracking *E. lenta* and ΦKL11 dynamics over 6 days in BALB/c mice (**Fig. S1 and Table S3B**). Together, these results demonstrate that *E. lenta* can evade ΦKL11-mediated killing within the gastrointestinal environment, even while maintaining high-levels of phage in the distal gut.

We hypothesized that the lack of competition with other bacterial strains and species could explain the ability of *E. lenta* DSM2243 to tolerate ΦKL11. To test this, we co-colonized GF mice with sensitive (DSM2243) and resistant (DSM15644) *E. lenta* strains (**Figs. S2A-D and Table S3C**), prior to ΦKL11 exposure in their drinking water for 2 weeks. Surprisingly, the abundance of both strains were unaffected by ΦKL11 (**Figs. 1H and S2E**). Finally, we repeated this experiment using a 4-member synthetic microbiota consisting of bacterial strains from the 4 most abundant phyla in the human gut: *E. lenta* DSM2243, *Bacteroides thetaiotaomicron* VPI-5482, *Escherichia coli* MG1655, and *Clostridium innocuum* DSM1286 (**Table S3D**). Yet again, *E. lenta* levels were not significantly altered by ΦKL11 (**Fig. 1I**). The other 3 community members were also stable over time (**Fig. S3**). These results highlight the reproducibility of *E. lenta* bacteriophage evasion independent of microbial community complexity or intraspecies competition.

### Bacteriophage resistance exists prior to phage exposure and is reversible

To more directly assess the potential emergence of bacteriophage resistant cells, we recovered a mixed population of *E. lenta* from endpoint stool samples collected from monocolonized mice treated with active or heat-killed (HK) ΦKL11 (**Fig. 1D-G**). *E. lenta* growth in these initial stool-derived *in vitro* communities was comparable between groups (0.73±0.08 active vs. 0.81±0.09 HK, *p*-value=0.28, Mann-Whitney U test). However, ΦKL11 was only able to form plaques on *E. lenta* recovered from mice treated with HK ΦKL11 (**Figs. 2A,B**). Bacteriophage resistance was stable following three additional passages in rich media (**Figs. 2C,D**), consistent with stable maintenance of ΦKL11 levels in these cultures (**Figs. 2E,F**). This effect was replicated *in vitro*, with bacteriophage resistance detectable following exposure of *E. lenta* DSM2243 to a low multiplicity of infection (MOI) of ΦKL11 (**Fig. S4**).

**Figure 2.**
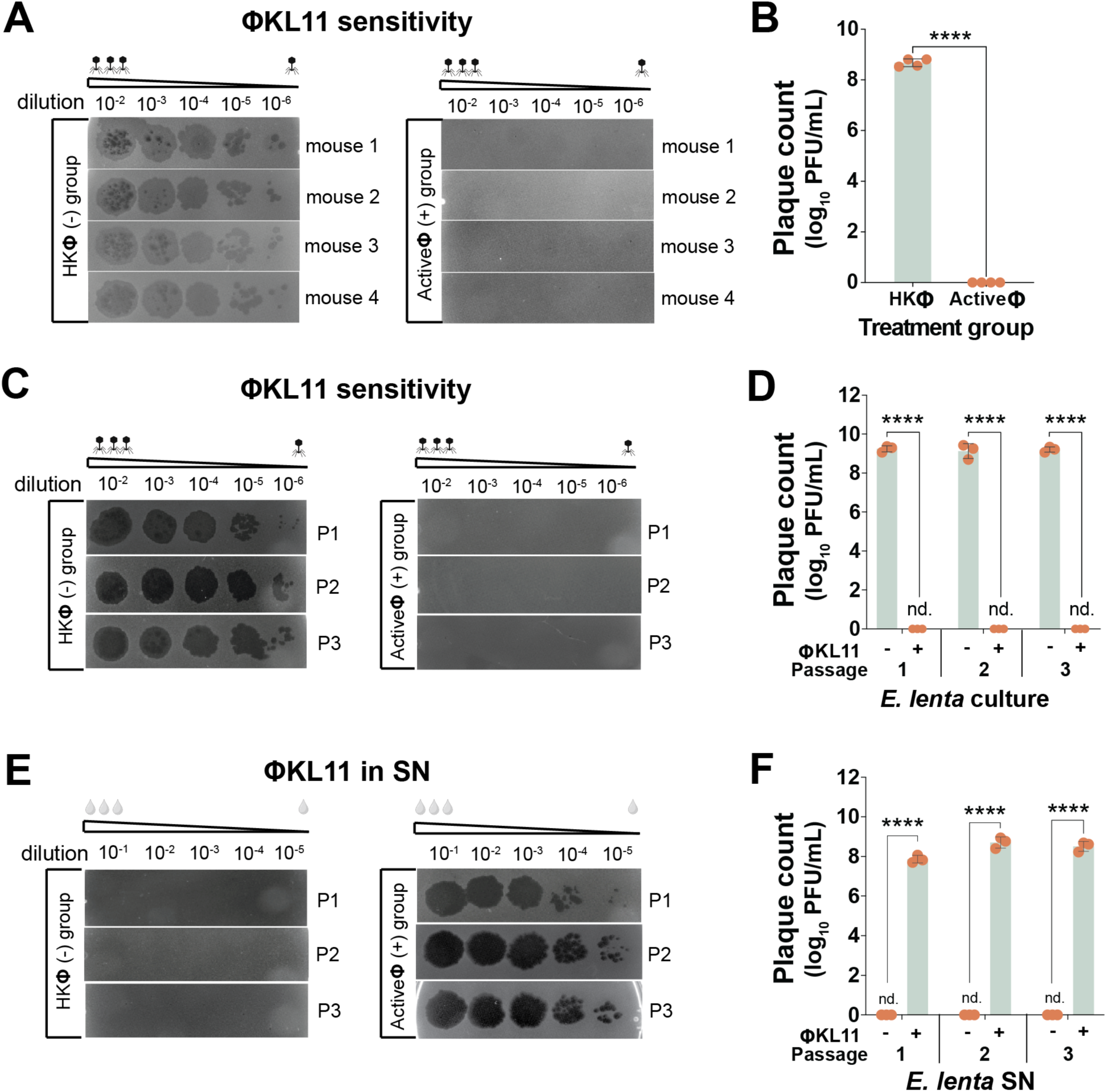
The *E. lenta* population becomes phage resistant upon ΦKL11 exposure in mice. **(A,B)** *E. lenta* recovered from the stool of monoassociated mice (Figures 1D-G) treated with active ΦKL11 are resistant to plaque formation relative to heat-killed (HK) controls. **(C,D)** The phage resistance phenotype is stable following three sequential passages in liquid media for a representative mouse-derived culture from each group. **(E,F)** ΦKL11 continues to be produced by passaged *E. lenta* cultures from mice treated with active ΦKL11, as indicated by plaque formation from culture supernatants (SN). **(A,C,E)** Images are representative of 2 technical replicates. **(B,D,F)** Values are mean±sem, dots represent the mean of each pair of technical replicates (n=3-4 biological replicates/group; \*\*\*\**p*<0.0001, Mann–Whitney test).

The rapid emergence of bacteriophage resistance in both mice and cell culture suggested the presence of pre-existing heterogeneity within *E. lenta* DSM2243. To quantify the frequency of Φ^S^ and Φ^R^ cells in ΦKL11-naive cultures, we sorted and analyzed single cells of *E. lenta* DSM2243 followed by plaque assays on solid media (**Fig. S5A**). Consistent with the overall Φ^S^ phenotype (**Figs. 1A-B**), 97.4% (37/38) of cells were Φ^S^ (**Fig. 3**); however, we did recover a single cell that was fully resistant to ΦKL11. We performed three subsequent passages for this Φ^R^ isolate (Φ^R1^) and a randomly selected sensitive isolate (Φ^S16^) followed by an additional round of single-cell sorting and plaque assays (**Fig. 3**). Despite the lack of selective pressure from ΦKL11, 29.2% (21/72) of the recovered cells from Φ^S16^ were now bacteriophage resistant. In contrast, Φ^R1^ had nearly entirely reverted to the sensitive phenotype, with 96.2% (75/78) of the recovered cells bacteriophage sensitive (**Fig. 3**). We also observed a broad range of plaque morphologies for the Φ^R1^ progeny, indicative of substantial phenotypic heterogeneity during outgrowth of each single cell (**Fig. S5B**). The reversible emergence of ΦKL11 resistance in the absence of selective pressure is consistent with phase variation, in which bacterial populations undergo stochastic, heritable and reversible switching between distinct phenotypic states^21,35^. This observation prompted us to determine whether ΦKL11 sensitivity and resistance correspond to distinct, reversible genomic configurations by performing long-read sequencing of distinct *E. lenta* subcultures.

**Figure 3.**
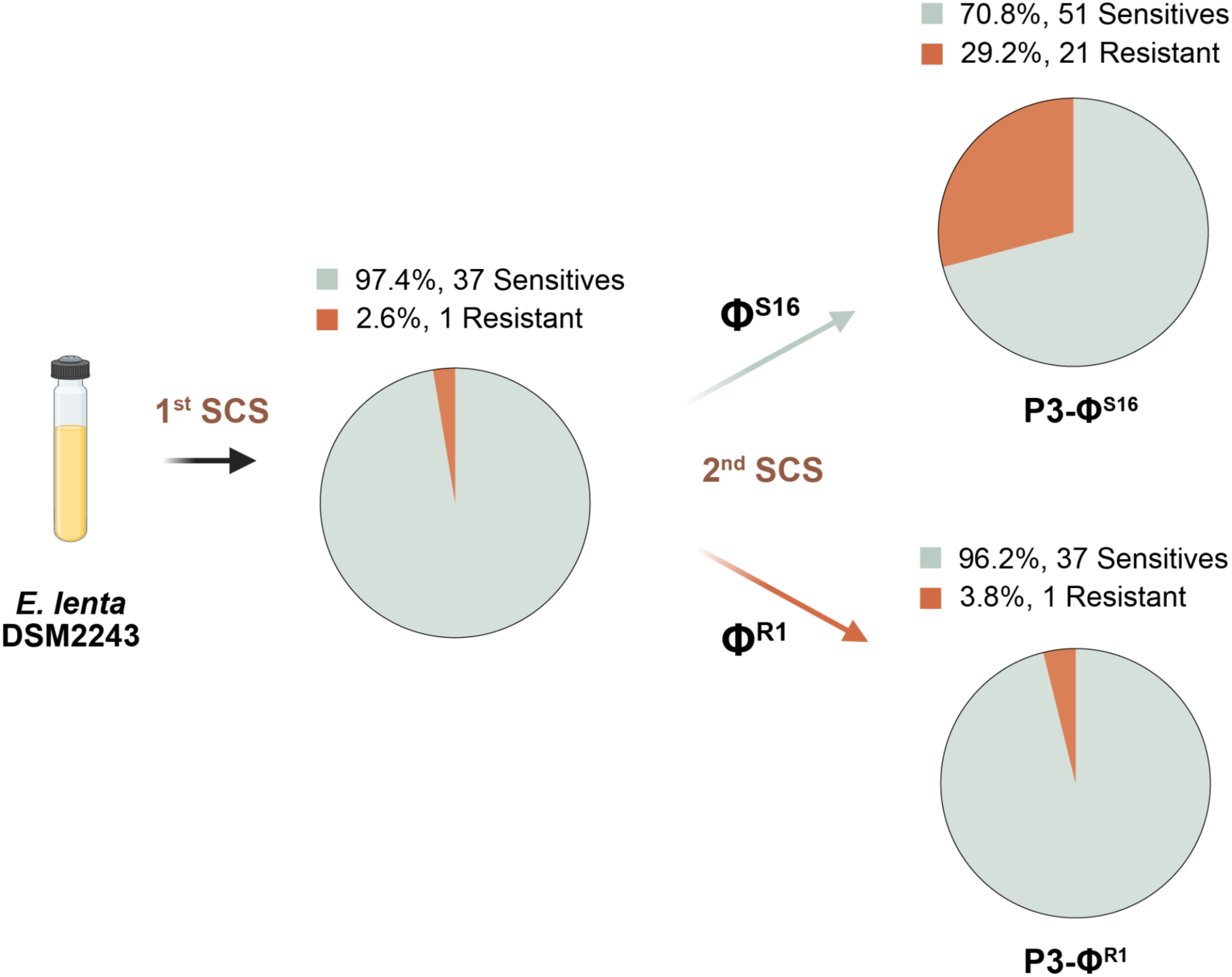
Single-cell sorting reveals that phage resistance exists prior to phage exposure and is reversible. Phage sensitivity by plaque assay after single-cell sorting (SCS) and recovery of 38 *E. lenta* DSM2243 cells (97.4% Φ^S^). We passaged a representative Φ^S^ and Φ^R^ isolate three times in BHI^A^ media prior to a second round of SCS, demonstrating the reversibility of this phenotype.

### Large genomic inversions occur in bacteriophage resistant *E. lenta*

We turned to whole-genome sequencing to evaluate the genetic basis for the observed phenotypic heterogeneity in bacteriophage sensitivity. Remarkably, we identified three distinct genomic structural variations (denoted SV0, SV1, and SV2) in the consensus genomes generated from long-read sequencing of independent *E. lenta* DSM2243 subcultures in the absence of phage selection (**Figs. 4A,B and Tables S4,S5**). SV1 was characterized by a 2.06 Mbp inversion (**Fig. 4A**) relative to SV0. SV2 had an additional 32.5 Kbp inversion relative to SV1 (**Fig. 4B**). We designed unique primer pairs for each structural variant (SV) to enable PCR-based identification of individual SVs (**Table S6**). Sanger sequencing was performed to validate the inversion junction sequences (**Table S5**).

**Figure 4.**
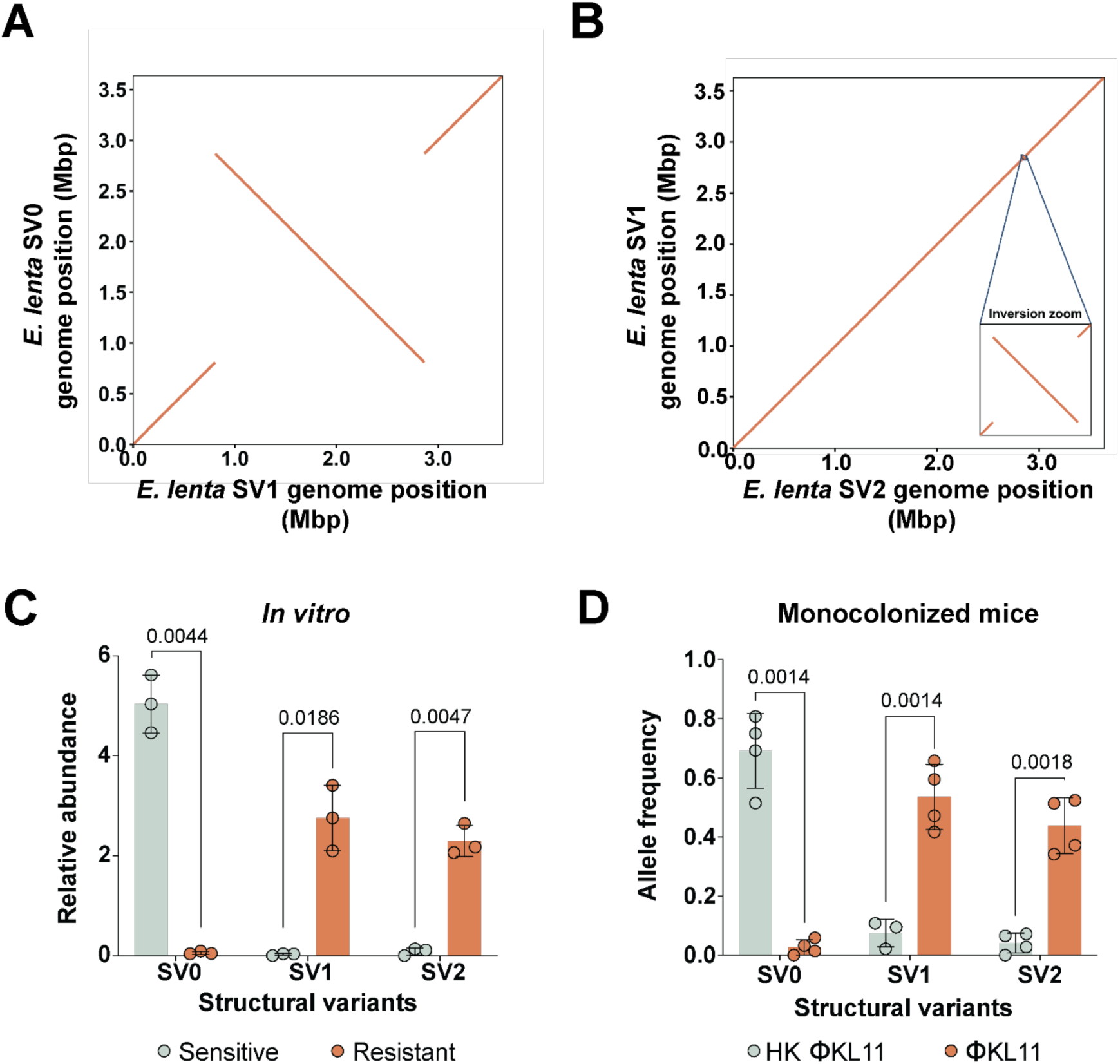
Large genomic inversions are selected during phage exposure in *E. lenta*. **(A,B)** Whole-genome alignment of *E. lenta* reference genomes reveals three unique SVs (SV0, SV1, and SV2). The y-axis shows reference genome coordinates, and the x-axis shows query coordinates (Mbp); lines denote collinear blocks and their orientation. **(A)** SV0 vs. SV1. **(B)** SV1 vs. SV2. **(C)** qPCR analysis of SV abundance in ΦKL11^R^ and ΦKL11^S^ isolates. Each dot represents an individual culture (**Figure S4**). **(D)** SV abundance based on long-read sequencing in *E. lenta* populations recovered from monoassociated mice. Each dot represents a mouse sample (Figure 2A**,B**). **(C,D)** Values are mean±sem (*p*-values, Student’s *t* tests).

To evaluate the generalizability of these large-scale chromosomal inversions within the *E. lenta* species, we analyzed an additional 10 strains using long-read sequencing (**Table S4**). Three *E. lenta* strains (539-5C, FCC8, and 943-4) exhibited large chromosomal inversions comparable to those observed in *E. lenta* DSM2243, spanning approximately 1.9–2.0 Mbp (**Table S5; Fig. S6A**). Sequence analysis of inversion junctions across DSM2243, 539-5C, FCC8, and 943-4 revealed that all large chromosomal inversions are flanked by an identical 16-bp inverted repeat (IR) sequence (5′-GATAAGCGCGGTTATG-3′; **Fig. S7**). In DSM2243, this IR defines the SV1 and SV2 boundaries, whereas in the other three strains it flanks the ∼2 Mbp inversion, supporting a conserved site-specific recombination mechanism^36^. In contrast, the smaller 32.5 Kbp inversion that was only detected in *E. lenta* DSM2243 has multiple features indicative of a transposon-associated event, including a flanking 39-bp IR (5′-TAAGCGCGGTTATGTAAAGTCGAGCGCNAGGGTCGGGGA-3′) and a centrally located putative IS1595-family transposase (ELEN_RS15785).

Next we sought to quantify the genomic structure of *E. lenta* in response to ΦKL11 infection. We performed an *in vitro* co-evolution experiment between *E. lenta* DSM2243 and ΦKL11 to select for phage-resistant populations. SV-specific PCR analysis of *in vitro* derived Φ^S^ and Φ^R^ *E. lenta* cultures revealed that SV0 was significantly enriched in Φ^S^ cultures, whereas SV1 and SV2 were enriched in Φ^R^ *E. lenta* (**Fig. 4C**). Consistent with these results, SV0 was significantly enriched in *E. lenta* cells recovered from mice treated with HK ΦKL11 and SV1/SV2 were significantly enriched in response to active bacteriophage (**Fig. 4D**). This data implicates the large 2.06 Mbp inversion in bacteriophage resistance, prompting us to further explore the genes at each inversion junction.

### Phage resistance is linked to differential expression of CPS clusters flanking each inversion

Both the large and small inversions detected in *E. lenta* DSM2243 re-arrange three gene clusters predicted to contribute to capsular polysaccharide synthesis (CPS; **Fig. 5A and Table S7**). Similarly, the single inversions found in the three additional *E. lenta* strains generated distinct SVs that rearranged a single capsular polysaccharide (CPS) gene cluster (**Fig. S6B**). Across all strains examined, these inversions repositioned CPS genes relative to a LuxR-like transcriptional regulator and/or the antitermination factor loaP. In contrast to DSM2243, where the inversions encompass three CPS gene clusters, the inversions in these strains predominantly involved a single CPS cluster.

**Figure 5.**
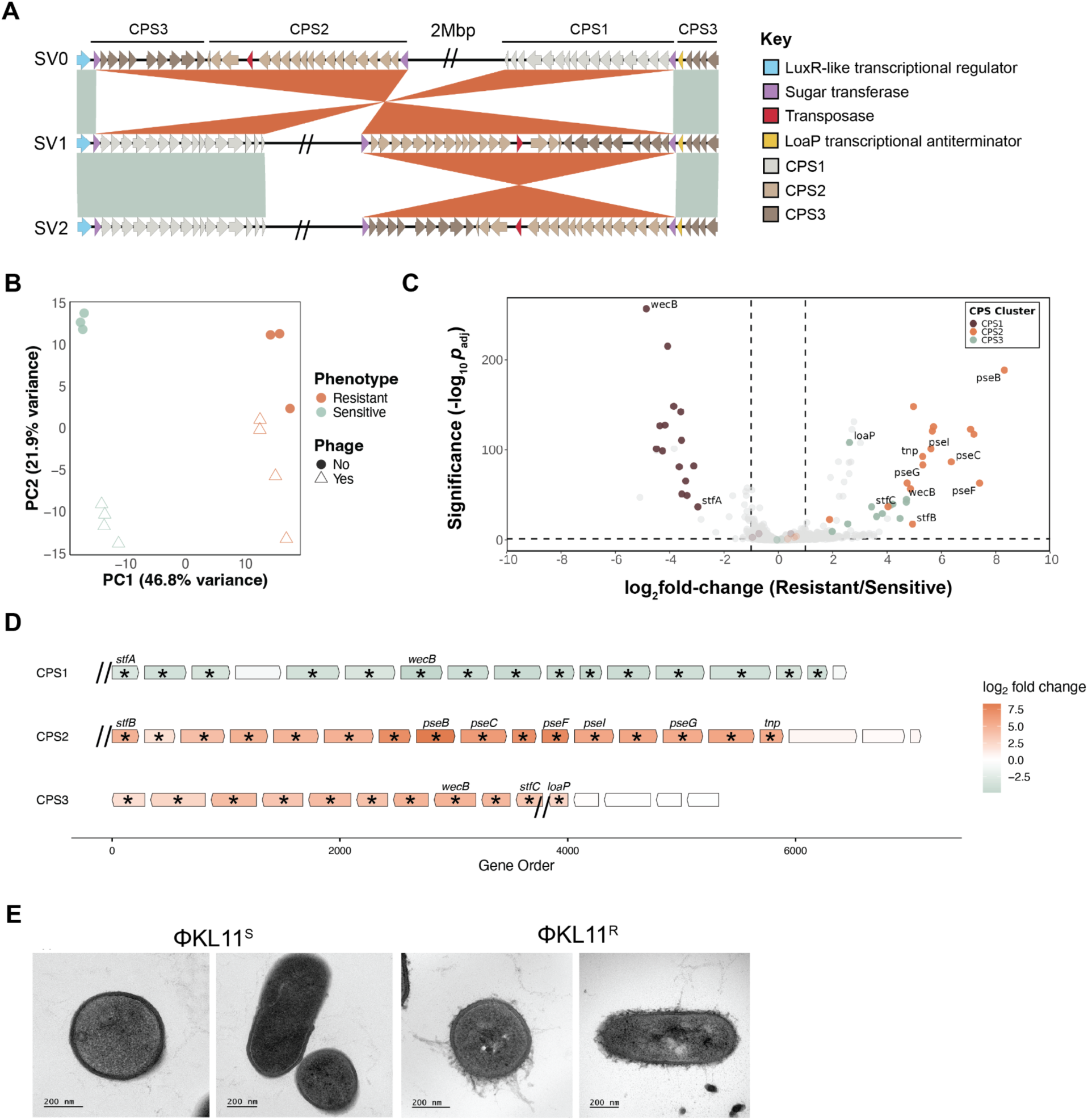
Phage resistance is linked to differential expression of CPS clusters flanking the large genomic inversion. **(A)** Comparative alignment of the flanking genomic neighborhoods of the three inversions identified in *E. lenta* (Figures 4A**,B**). Orange ribbons denote homologous blocks (crossed ribbons indicate inversions), and mint shading marks syntenic regions in the same orientation. Arrowed boxes represent CDS, highlighting key features of each CPS gene cluster (LuxR-like regulator, sugar transferases, transposase, LoaP). Double slashes (“//”) denote a 2-Mbp region omitted in SV1 and SV2. **(B)** Principal component analysis of *E. lenta* transcriptomes, colored by phenotype (orange: ΦKL11ᴿ; mint: ΦKL11^S^) and shaped by phage exposure (circles: untreated; triangles: ΦKL11-treated). **(C)** DEGs comparing ΦKL11^R^ and ΦKL11^S^ *E. lenta*. CPS genes are highlighted (CPS1: brown; CPS2: orange; CPS3: mint). Vertical dashed lines mark |log₂FC| = 1; the horizontal dashed line marks *p*_adj_≤0.05. **(D)** Zoomed-in view of each CPS gene cluster. Arrows indicate genes and transcriptional orientation, colored by log₂ fold change in ΦKL11^R^ vs. ΦKL11^S^ (orange: ΦKL11^R^ enriched; mint: ΦKL11^S^ enriched). Asterisks denote significantly differentially expressed genes; hash marks indicate inversion junctions. **(E)** Representative TEM images of ΦKL11-sensitive ΦKL11^S^ (left two panels) and ΦKL11^R^ (right two panels) cells. Scale bars, 200 nm.

We hypothesized that inversion of the three CPS gene clusters in *E. lenta* DSM2243 would alter their expression, resulting in differences in the bacterial cell surface that enable bacteriophage resistance. To test this hypothesis, we used RNA sequencing (RNA-seq) to compare the transcriptomes of Φ^S^ and Φ^R^ *E. lenta* DSM2243 in the presence or absence of ΦKL11. Bacterial gene expression was markedly distinct with respect to bacteriophage sensitivity (**Fig. 5B**), including 226 differentially expressed genes (DEGs; **Fig. 5C and Table S8**). Notably, a total of 42 DEGs were found within the identified CPS gene clusters (**Figs. 5D and Table S8**), including a clear increase in the expression level of CPS genes found on one side of the large inversion (denoted as CPS2/CPS3) at the expense of the other side (CPS1). The phenotypic consequences of these transcriptional changes were validated by transmission electron microscopy. We observed a marked difference in the cell surface of Φ^S^ and Φ^R^ *E. lenta* (**Fig. 5E**), with Φ^S^ cultures displaying an ordered capsule in contrast to the more extensive nature of the Φ^R^ capsule.

We also leveraged our RNA-seq data to analyze gene expression responses to ΦKL11 infection. This analysis identified a total of 446 differentially expressed genes (DEGs), of which 57 were shared between Φ^S^ and Φ^R^ *E. lenta*, while 327 and 62 DEGs were uniquely associated with Φ^S^ and Φ^R^ hosts, respectively (**Fig. S8A**). Notably, four genes in CPS2/CPS3 were significantly upregulated exclusively in the phage-sensitive host (**Figs. S8B-E**). In contrast, the CPS2 gene *pseG* (UDP-2,4-diacetamido-2,4,6-trideoxy-beta-L-altropyranose hydrolase) was significantly downregulated only in the phage-sensitive host upon ΦKL11 exposure (**Fig. S8F**). Although *pseG* resides within the CPS2 locus, the divergent regulation of *stfB* (sugar transferase) and *pseG* upon ΦKL11 exposure suggests modular control of capsule biosynthesis. Additionally, 7.91% of the genes upregulated upon phage infection exclusively in the phage-sensitive host correspond to insertion sequence (IS) elements, primarily IS3- and IS256-family transposases (**Fig. S8G and Table S9**), consistent with recent reports of phage-induced IS256 transposition in E. faecalis^37^.

### Bacteriophage ΦKL11 has a broad host range

To assess the host range of ΦKL11, we screened 107 *E. lenta* strains for their ability to form plaques upon infection with a range of phage titers on solid media and quantified each strain’s susceptibility to ΦKL11 infection. We found that 50.46% (54/107) of the tested *E. lenta* strains could form plaques, indicating sensitivity to ΦKL11. In contrast, no plaques were detected for the remaining strains, even at the highest phage titers tested, suggesting that these strains are not susceptible to ΦKL11 (**Fig. 6A**). Bacterial cell density (OD₆₀₀) was comparable between groups (0.93 ± 0.2 host vs. 0.89 ± 0.21 non-host, *p*-value=0.52, Mann-Whitney U test). Plaque-forming strains produced a mean titer of 8.57 ± 0.84 log₁₀ PFU/mL, spanning a broad range from 6.17 to 9.62 log₁₀ PFU/mL (**Fig. 6B**), with some strains comparable to the type strain DSM2243 (9.629 ± 0.051 log₁₀ PFU/mL). Mapping ΦKL11 susceptibility onto the core genome phylogeny of *E. lenta* revealed that host and non-host strains are broadly interspersed across the tree (**Fig. S9**), indicating that phage sensitivity does not strictly follow phylogenetic relatedness and is instead likely determined by specific genetic features that vary among closely related strains.

**Figure 6.**
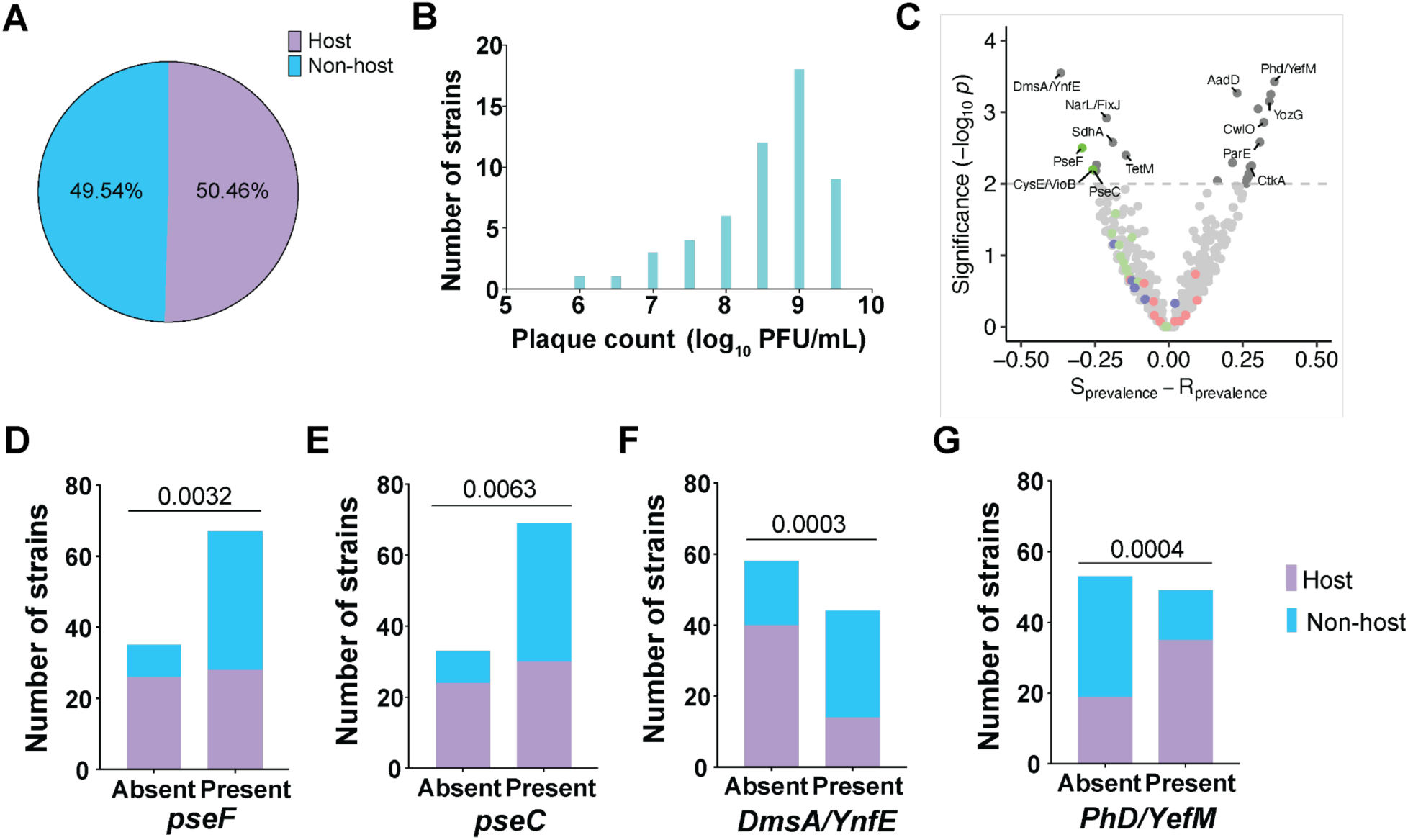
Bacteriophage ΦKL11 has a broad host range. **(A)** ΦKL11 host range analysis based on plaque assay across 107 *E. lenta* strains: non-host (cyan) or host (lavender). **(B)** Histogram of ΦKL11 titers across susceptible *E. lenta* strains, shown as log₁₀ PFU/mL; bars indicate the number of strains per titer bin. **(C)** Genome-wide association of orthologous group presence/absence with ΦKL11 susceptibility phenotype (n=102 strains with genome sequences available; **Table S11**). Each point is an orthologous group; the x-axis shows the difference in prevalence between host and non-host isolates, and the y-axis shows −log₁₀ *p*-value from Fisher’s exact tests. The dashed line marks the significance threshold (−log₁₀ *p*=2). Labeled points denote representative genes. **(D-G)** Counts of sensitive and resistant isolates stratified by gene presence or absence for four loci (*pseF*, *pseC*, *dmsA*/*ynfE*, *phd*/*yefM*). *p*-values, Fisher’s exact tests.

To identify genetic features distinguishing phage host and non-host strains, we conducted a genome-wide presence–absence association analysis by comparing orthologous gene groups across 102 *E. lenta* isolates (**Tables S10,S11**). Using Fisher’s exact test, we identified genes whose presence was significantly correlated with ΦKL11 sensitivity or resistance. In total, 11 genes were significantly enriched in non-host strains, and 16 genes were significantly enriched in host strains (*p*<0.01, Fisher’s exact test; **Fig. 6C**). Notably, two genes within CPS2, *pseF* (pseudaminic acid cytidylyltransferase) (**Fig. 6D**) and *pseC* (UDP-4-amino-4,6-dideoxy-N-acetyl-beta-L-altrosamine transaminase) (**Fig. 6E**) were more common in non-host strains, consistent with the ability of altered cell surface polysaccharides to evade bacteriophage infection. The top gene enriched in non-host strains was dimethyl sulfoxide reductase subunit (**Fig. 6F**, *dmsA/ynfE*), a molybdenum-dependent enzyme associated with oxidative stress and bacterial pathogenesis^38^. The top gene enriched in host strains was the antitoxin component (*phd/yefM*) of a type II toxin–antitoxin module implicated in bacterial stress responses^39^ (**Fig. 6G**).

Finally, we examined the distribution of capsule polysaccharide (CPS) gene clusters across 102 *E. lenta* genomes. Gene presence/absence based on eggNOG ortholog annotations revealed substantial variability in CPS loci across strains (**Fig. S10A**). To quantify the prevalence of complete CPS clusters, we defined cluster presence using an 85% sequence similarity cutoff for CPS genes (see **STAR Methods**). Using this criterion, CPS1 was detected in 34.95% of genomes, CPS2 in 25.24%, and CPS3 in 32.04% of genomes (**Fig. S10B**). We then asked whether the presence of these CPS clusters was associated with ΦKL11 susceptibility. Comparison of host and non-host strains showed no significant association for CPS1 or CPS3 (**Fig. S10C**). In contrast, CPS2 was significantly more prevalent among ΦKL11 non-host strains, supporting a broader role of the CPS2 gene cluster in protecting from phage infection across multiple *E. lenta* strains.

## DISCUSSION

This work represents the first in-depth experimental characterization of a bacteriophage that targets the prevalent and disease-associated gut Actinomycetota *Eggerthella lenta.* ΦKL11 exhibits robust lytic activity *in vitro* and a broad host range, resulting in plaque formation on >50% of the tested strains despite considerable strain-level variation in gene content^22,40^. Thus, ΦKL11 represents a valuable and much-needed new tool for understanding *E. lenta* biology, especially given the recent studies highlighting the broad impacts of *E. lenta* on gut microbial ecology^41,42^, the metabolism of dietary^26^ and pharmaceutical^23,24,28^ small molecules, and immune system function^30,43^.

Despite the clear ability of ΦKL11 to lyse *E. lenta* during growth on solid and in liquid media, our experiments in gnotobiotic mice did not reveal any detectable changes in *E. lenta* colonization levels. The lack of any transient decrease in colonization supports our working model that pre-existing heterogeneity in *E. lenta* genome structure, gene expression, and cell surface phenotype enable robust evasion of bacteriophage targeting. Interestingly, our results were not affected by the presence of other gut bacterial strains or species, emphasizing how variability within a bacterial strain can matter more than overall gut microbial community structure. These results also highlight the critical need to move beyond simple cell culture experiments to incorporate mice and/or other more physiologically relevant models. Notably, a simple refinement to our *in vitro* experiments to include multiple rounds of sequential phage infection amplified and validated the ability of *E. lenta* to evade bacteriophage infection. Continued modifications of these cell culture assays to incorporate additional aspects of the gastrointestinal tract (e.g., host cells, mucus, cytokines) could further accelerate and improve the translational relevance of these findings and provide new opportunities for isolating additional bacteriophage that more robustly target *E. lenta* within the gastrointestinal tract.

Repeated isolation efforts from the same wastewater treatment plant in San Francisco led to the isolation of closely related bacteriophage in two subsequent years, indicating that sampling more geographically distinct locations could well support the isolation of a more diverse set of phages that target *E. lenta*. Notably, the only previously published isolation of a bacteriophage that targets *E. lenta* was from a wastewater treatment plant in Kiel, Germany. This study revealed another distantly related siphovirus^44^; BLASTn alignment with ΦKL11 revealed no detectable sequence homology. CRISPR spacer matching^19^ and prophage analysis^45^ indicate that the diversity of bacteriophages targeting *E. lenta* extend well beyond the isolates described here. These include small dsDNA tailed phages detected in human gut metagenomes, additional uncultivated phage lineages inferred from CRISPR spacer matches, and more than ten distinct temperate phage clades integrated within *E. lenta* genomes.

Our results also indicate a potential tradeoff between bacteriophage evasion and interactions with the mammalian host. The default structural variant for *E. lenta* DSM2243 within the mouse gastrointestinal tract was SV0, which is indicative of increased sensitivity to phage infection. Exposure to ΦKL11 led to a marked shift in *E. lenta* population structure to favor the structural variants associated with bacteriophage resistance, which was associated with altered capsular polysaccharide biosynthesis. These changes to the *E. lenta* capsule likely have important implications for host immunity, given the key role of capsular polysaccharides for immune activation in other gut bacterial species^46,47^ and the recently described role of select *E. lenta* strains in driving intestinal immune activation and colitis^30^. Notably, the CPS2 gene cluster encodes homologs of the pseudaminic acid biosynthesis pathway, which has been implicated in immune evasion in diverse pathogenic bacteria, including *Acinetobacter baumannii, Campylobacter jejuni,* and *Helicobacter pylori*^48–50^. Our transcriptomic and comparative genomic analyses further implicated this pathway in bacteriophage evasion and ΦKL11 host range, emphasizing how bacteriophage targeting can change the expression of bacterial pathways implicated in virulence.

One of the most striking aspects of the mechanism for bacteriophage evasion described herein is the physical size of the inversions observed, leading to changes in genome structure on the megabase scale, exceeding 50% of the total genome size in *E. lenta* DSM2243. This observation is in stark contrast to the more well-studied but far shorter promoter inversions that control capsular polysaccharide biosynthesis and bacteriophage sensitivity in the model gut bacterium *Bacteroides thetaiotaomicron*^35^. A systematic analysis of DNA inversions in the gut microbiome revealed a median length of ∼100-200 bp^51^; however, longer inversions were likely missed due to read-length and bioinformatic limitations. We discovered >1 Mbp inversions in 4 distinct *E. lenta* strains, highlighting the value of re-sequencing multiple stocks of the same gut bacterial species to assemble closed genomes that permit the comprehensive analysis of inversions of any size. This remarkable degree of genomic plasticity emphasizes the need to move beyond simply cataloguing the taxonomic and gene copy number within host-associated microbial communities to assess differences in gene orientation at the single cell level. Furthermore, our work demonstrates that the megabase-scale rearrangements made possible by engineered recombinases^52,53^ are also found in nature.

The size of these inversions also raises intriguing questions as to the detailed molecular mechanisms responsible. We hypothesize that the *E. lenta* genome may physically fold in a manner that brings distant genomic regions into close proximity, which could be tested using emerging methods for analyzing the three dimensional structure of bacterial genomes^54^. Close inspection of the flanking regions of each inversion revealed a conserved 16-bp IR across all 4 *E. lenta* strains. In many bacterial systems, such IRs serve as recognition sites for nearby site-specific recombinases that catalyze DNA inversion^55–58^. However, we did not detect recombinase genes adjacent to the inversion breakpoints in *E. lenta*. Together, these observations suggest that the megabase-scale inversions may arise through recombination between the conserved IRs, potentially mediated by recombination machinery acting in trans^59^. In addition, our data suggests that the smaller 32-Kbp inversion (SV2) represents an active transposon, driven by the 39-bp IR and an IS1595-family transposase. Thus, two distinct mechanisms of inversion can result in nested SVs that vary between gut bacterial strains.

Another critical question is the mechanism responsible for the marked differences in gene expression between structural variants. Multiple observations support a more complicated model than a simple promoter swap. First, the structural variants arise from large chromosomes that relocate entire CPS gene clusters together with their surrounding genomic context, rather than simply inverting a small promoter region upstream of a single operon. Second, the transition from phage-sensitive to resistant states is accompanied by coordinated transcriptional changes across multiple CPS loci, with CPS1 expression decreasing while CPS2 and CPS3 increase. Finally, the inversions reposition CPS loci relative to nearby regulatory elements, including a LoaP transcriptional antiterminator and a LuxR-like regulator, suggesting that altered regulatory context may contribute to these expression differences. Together, these observations support a working model in which the large chromosomal inversion rewires capsule regulation by repositioning CPS loci relative to two opposing regulators. LoaP promotes transcription of the CPS locus with which it is linked^60^, whereas the LuxR-like regulator may modulate expression of adjacent CPS clusters. In this model, inversion toggles regulatory control between CPS1 and CPS2/3, producing alternative capsule states associated with phage sensitivity and resistance.

More broadly, these results add to the growing literature on bacteriophage evasion strategies across host-associated and environmental ecosystems. Bacteria have evolved diverse strategies to evade bacteriophage infection, including CRISPR-Cas systems, abortive infection, toxin-antitoxin modules, and restriction-modification systems^61^. However, despite encoding multiple defense systems^19,22,40^, our data indicates that phase variation in capsule production is the primary mechanism through which *E. lenta* evades ΦKL11. Phase variation represents a comparatively underexplored mechanism, involving reversible, stochastic changes in gene expression that generate phenotypic heterogeneity^62^. These changes can arise through mechanisms such as inversions (as in our system), slipped-strand mispairing during DNA replication, or epigenetic modifications such as DNA methylation of regulatory regions^62^. Phase variation is also critical for bacteriophage evasion in the well-studied gut bacterium *B. thetaiotaomicron*^35^, indicating that this may be a general strategy enabling the long-term co- existence of lytic bacteriophage and their host bacteria within the gut environment and prompting more in-depth studies of phase variation across more diverse habitats.

Finally, this study has clear implications for the emerging field of microbiome editing^12^. Consistent with work in model organisms^17,37,63^, we are far from achieving complete decolonization of *E. lenta* despite administering a high dose of lytic bacteriophage. While it is possible that engineered bacteriophage, phage cocktails, and/or co-administration of antibiotics or other selective factors will increase potency, these results also emphasize the challenges in overcoming the long-term co-evolution of bacteriophages with their gut bacterial hosts. On the other hand, the continued heterogeneity of *E. lenta* even during phage exposure suggests that there is always a subset of cells conducive to gene delivery by ΦKL11. With continued refinement of gene overexpression systems^28,33^, this property could enable stable delivery of individual genes, operons, or even entire biosynthetic pathways for therapeutic purposes.

## METHODS

### Key resources table

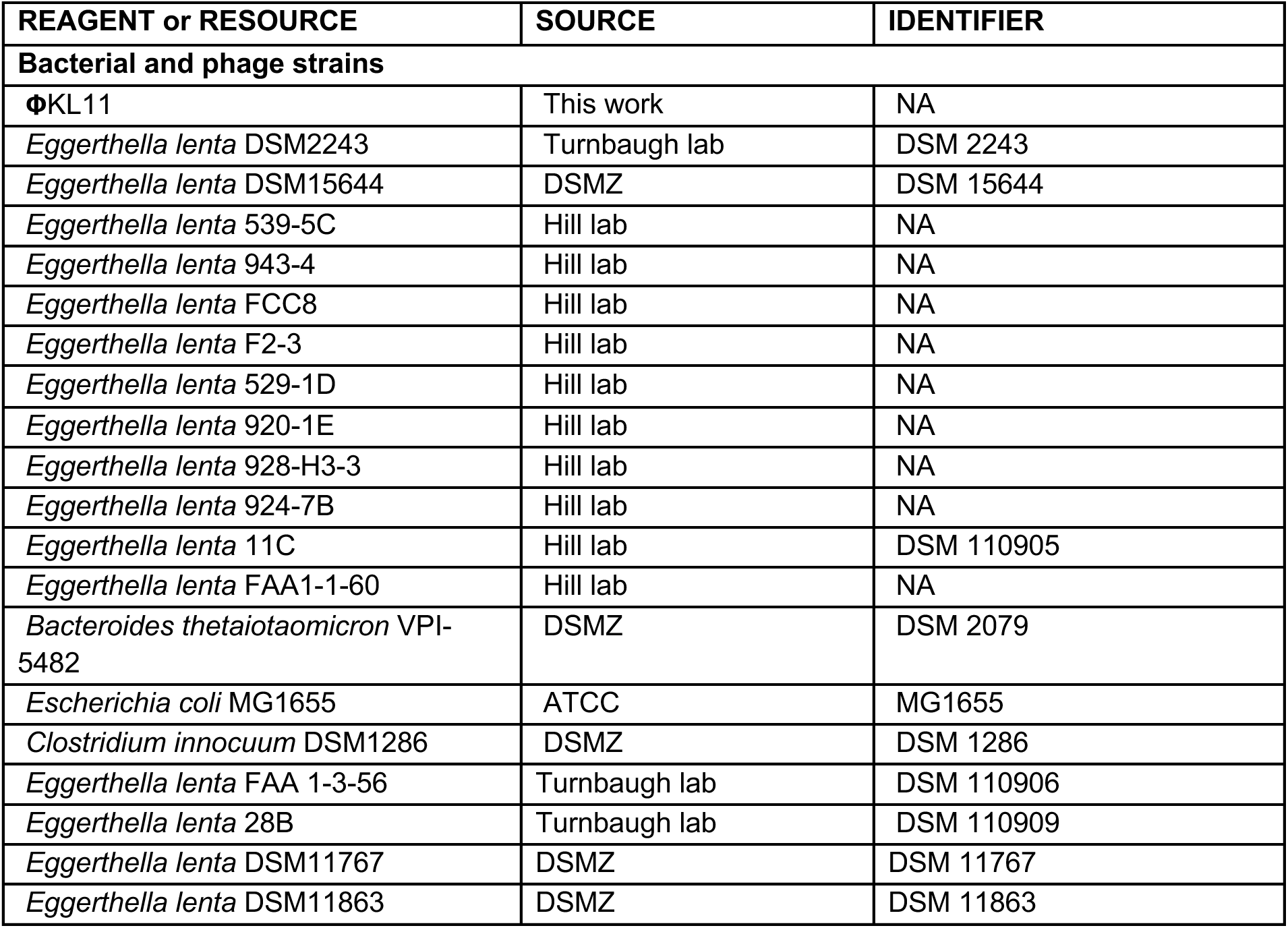
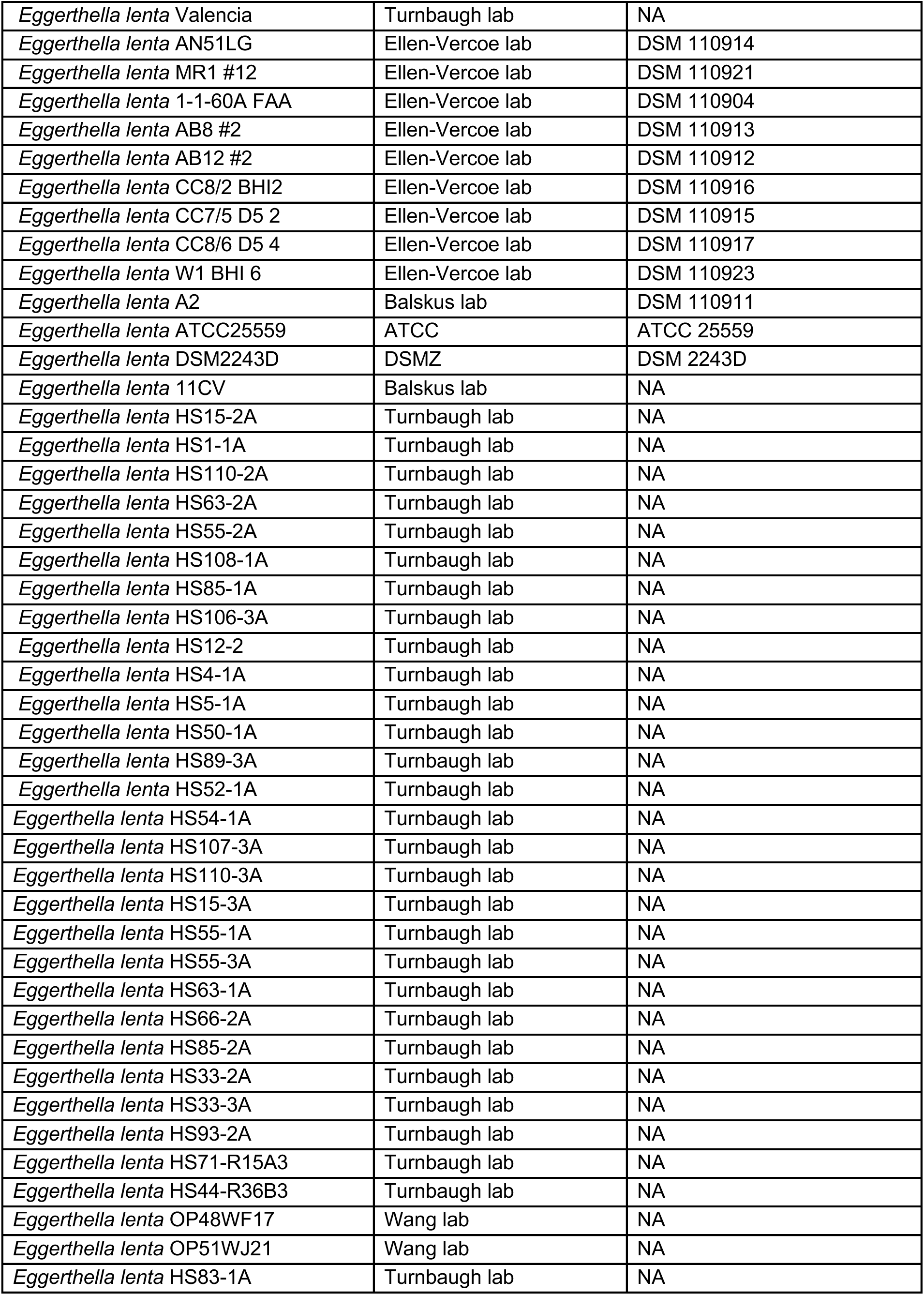
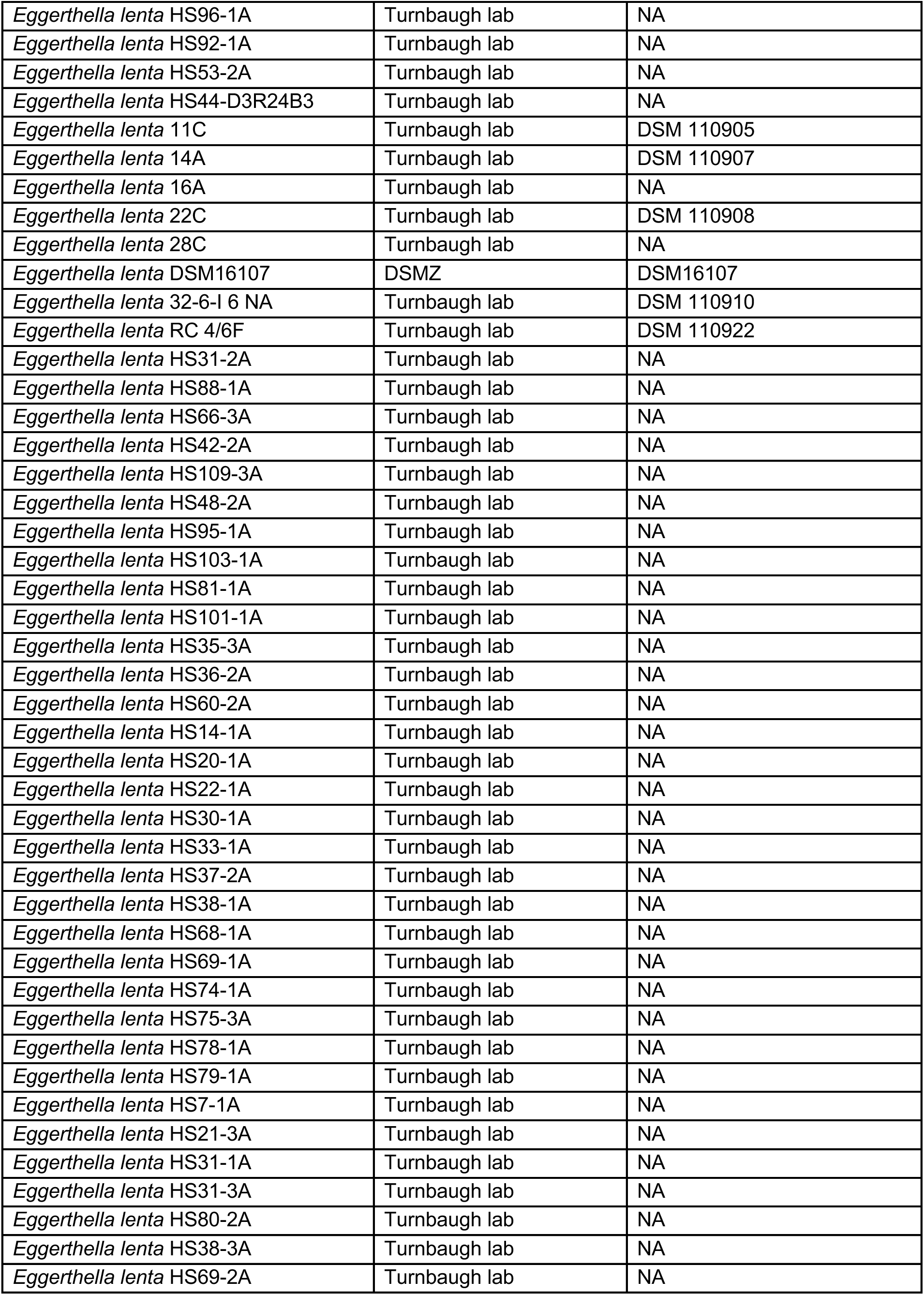
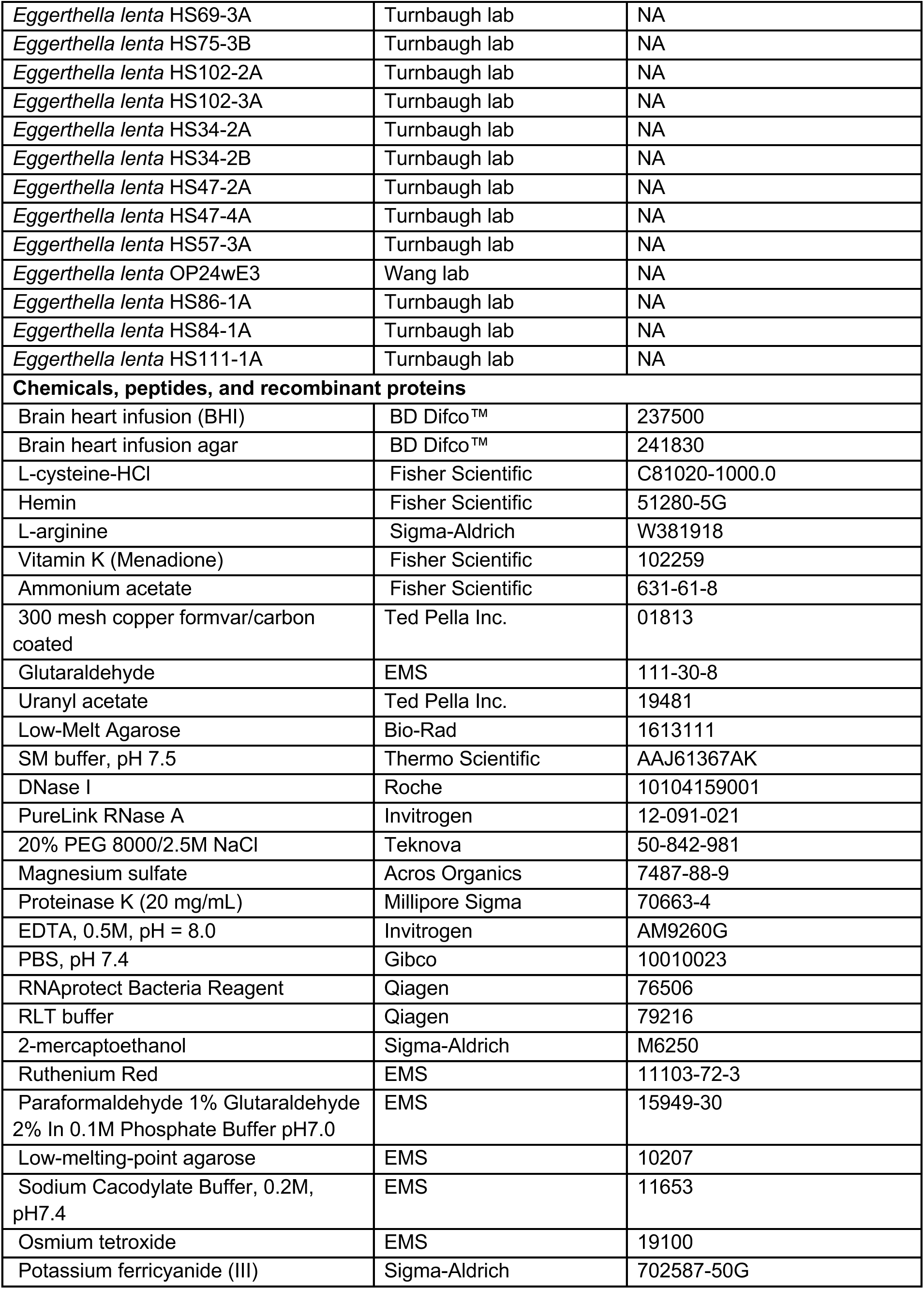
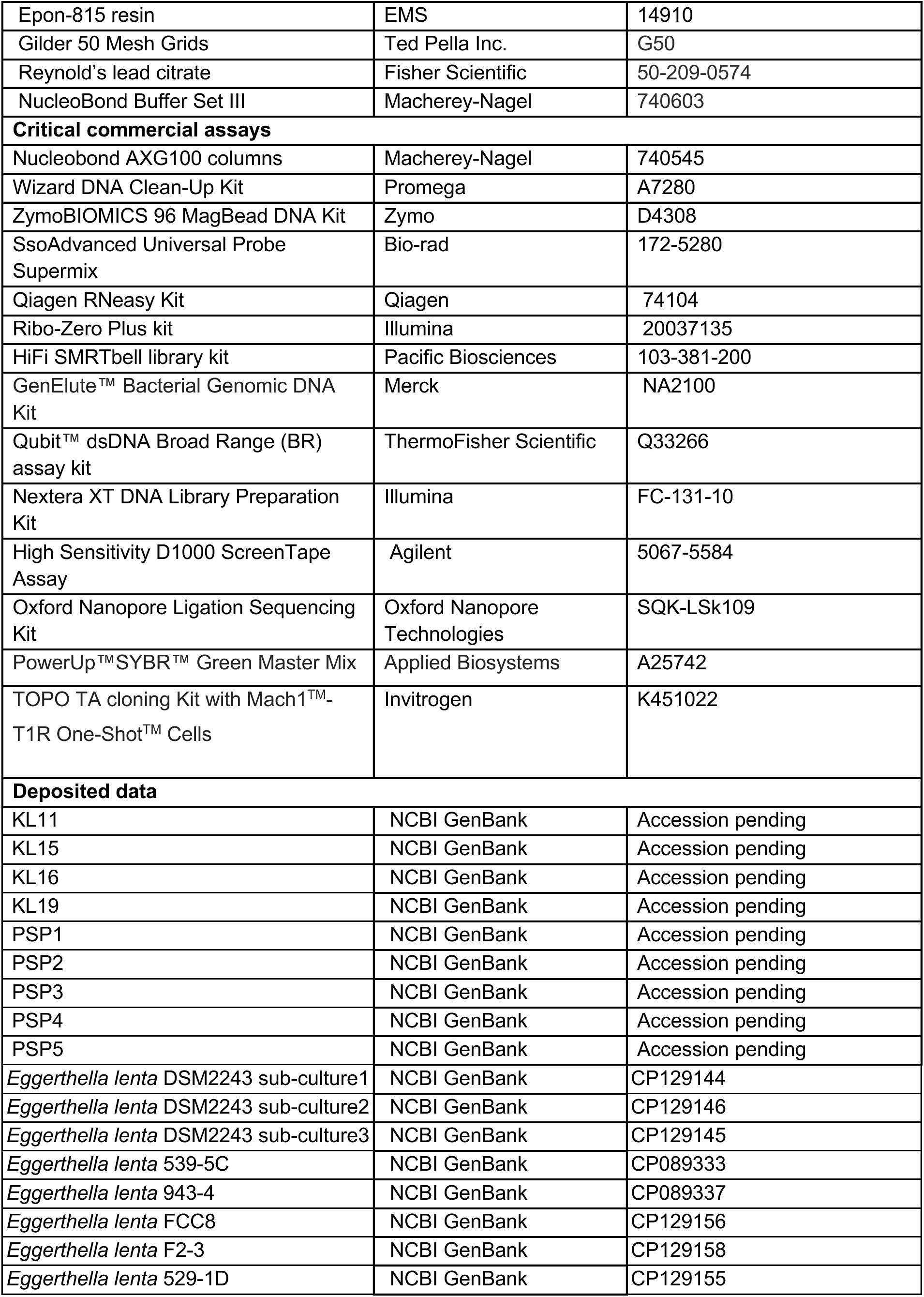
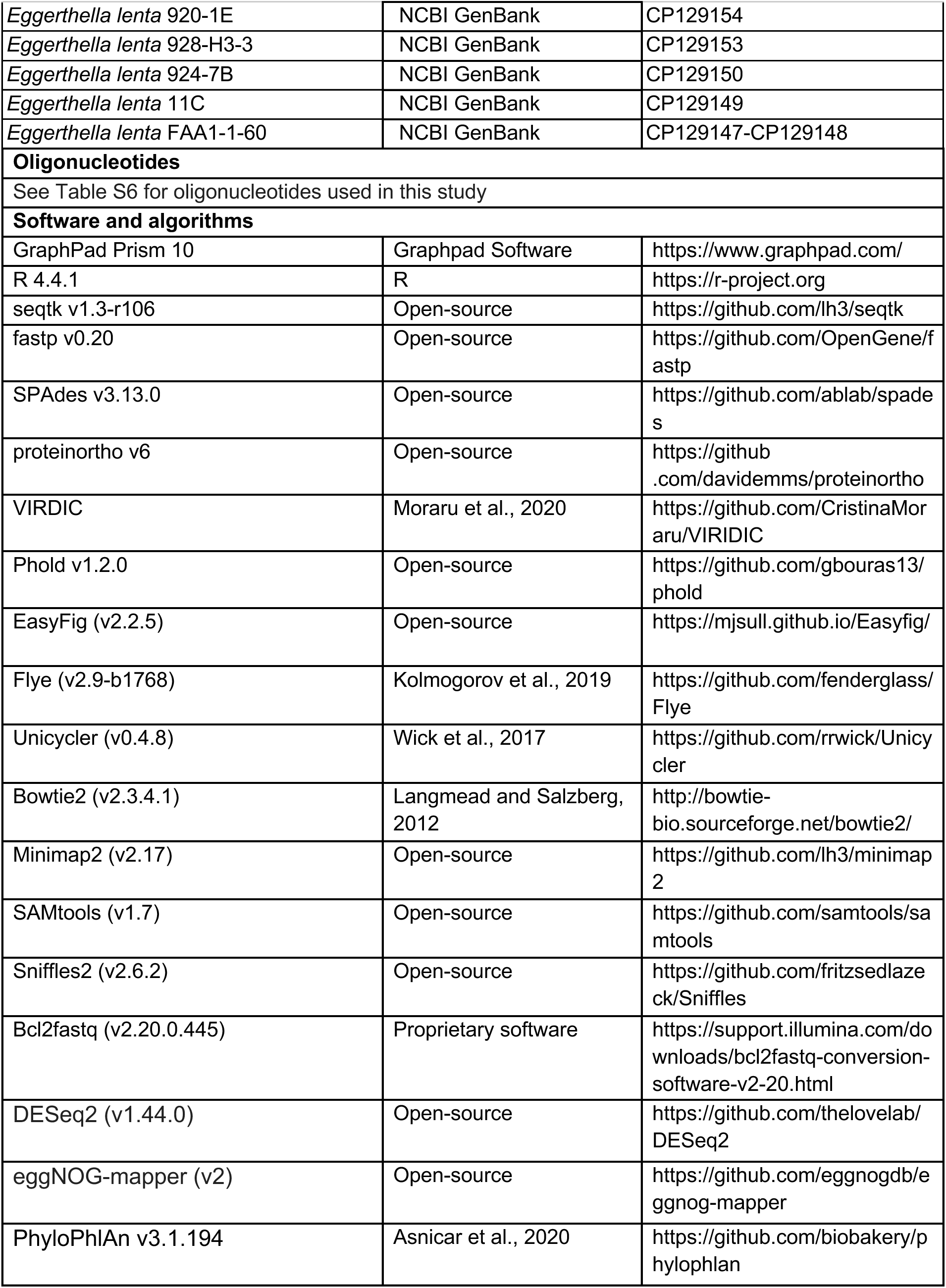

### Resource Availability

#### Lead Contact

Further information and requests for resources and reagents should be directed to and will be fulfilled by the Lead Contact, Peter Turnbaugh.

#### Material Availability

Newly isolated bacteriophages described in this study are available from the corresponding author upon reasonable request.

#### Data and Code Availability

The accession number for the sequencing data reported in this paper is NCBI Sequence Read Archive: BioProject Accessing Pending.

### Experimental Model and Subject Details

#### Mice

All animal experiments were conducted under protocols approved by the University of California San Francisco Institutional Animal Care and Use Committee. GF C57BL/6J and BALB/c mice were born and maintained in the UCSF Gnotobiotic Core Facility. Littermates were randomly assigned to experimental groups. All experiments were performed using wild-type C57BL/6J or BALB/c mice. C57BL/6J mice were 8 weeks of age at the start of the experiment, and BALB/c mice were 9–12 weeks of age. Germ-free experiments used either mixed-sex cohorts or male-only mice. Detailed information on all mice used in this study is provided in **Table S3**.

#### Bacteria culturing and growth curves

Bacterial strains were cultured at 37°C in an anaerobic chamber (Coy Laboratory Products) with an atmosphere composed of 2-3% H_2_, 20% CO_2_, and the balance N_2_. Culture media were based on brain heart infusion (BHI). Depending on the experiment, we used either BHI supplemented with 1% L-Arginine (BHI^A^) or BHI supplemented with L-Cysteine-HCl (0.05% w/v), hemin (5 μg/ml), 1% L-Arginine, and vitamin K (1 μg/ml) (BHI^CHAV^). All media were pre-equilibrated in the Coy chamber for at least 12 h prior to culturing.

Growth curves of *E. lenta* in the presence or absence of phage were performed in clear, flat-bottom 96-well polystyrene plates (non-tissue culture treated). BHI^CHAV^ broth and overnight *E. lenta* cultures were 100-fold diluted and added to each well to a starting inoculum of OD₆₀₀ = 0.003, followed by addition of phage in a 10-fold serial dilution. Plates were incubated at 37 °C under anaerobic conditions, and optical density at 600 nm (OD₆₀₀) was measured every 30 min for 48-72 h using a microtiter plate reader (BioTek Epoch 2, Agilent).

#### Bacteriophage isolation, purification, and propagation

Sewage samples were supplemented with 10× BHI^CHAV^ to achieve a final composition of 1× BHI^CHAV^, with wastewater comprising 90% (v/v) of the final volume. The medium was aliquoted into 3 mL portions in sterile tubes and pre-reduced for 12 h under anaerobic conditions. *E. lenta* DSM2243 was prepared by inoculating colonies from freshly streaked BHI^CHAV^ agar plates into BHI^CHAV^ broth and incubating anaerobically at 37 °C for 48 h. Enrichment was initiated by subculturing overnight cultures 1:50 (v/v) into pre-reduced wastewater–BHI^CHAV^ aliquots and incubating anaerobically at 37 °C for 16-24 h. Enrichment cultures were centrifuged at 12,000 × *g* for 2 min, and supernatants were sterile-filtered through 0.22 µm filters to obtain cell-free filtrates. For plaque assays, 1 mL of overnight *E. lenta* culture was pelleted (10,000 × *g*, 1 min), resuspended in 100 µL residual medium, and 100 µL of concentrated cells was mixed with 100 µL of filtrate. The mixture was combined with 5 mL molten 0.2% top agar and poured onto pre-reduced BHI^CHAV^ agar plates. Plates were incubated anaerobically at 37 °C and examined for plaque formation after 24-48 h incubation. Individual plaques were isolated and purified through three rounds of plaque picking and streaking. For *E. lenta* phage enumeration, 200 µL of *E. lenta* DSM2243 culture (48–72 h) was mixed with 200 µL of ΦKL11 phage stock (∼10^9^ PFU/mL) in a 15 mL conical tube and combined with 3 mL of molten 0.2% agarose (55°C). The mixture was overlaid onto BHI^CHAV^ agar plates and incubated anaerobically at 37 °C for 48 h. All *E. lenta* phages isolated from sewage samples are listed in **Table S1**.

#### Phage genomic DNA extraction, sequencing, and assembly

Phage genomic DNA (gDNA) was extracted following a modified Wizard DNA Clean-Up protocol developed by the Center for Phage Technology (Promega A7280). Briefly, 10 mL of phage lysate was treated with DNase I and RNase A, gently mixed, and incubated at 37°C for 30 min to remove contaminating host nucleic acids, following the manufacturer’s instructions. PEG/NaCl solution (20% PEG 8000, 2.5 M NaCl) was then added, and samples were incubated at 4°C overnight to precipitate phage particles. Lysates were centrifuged at 10,000 × *g* for 20 min at 4°C, and the supernatant was discarded. The pellet was resuspended in 5 mM MgSO_4_, transferred to a clean tube, and clarified by centrifugation to remove residual impurities. The resulting supernatant was transferred to a fresh tube, and Proteinase K and EDTA were added to final concentrations of 100 µg/mL and 10 mM, respectively, followed by incubation at 50°C for 30 min. Phage gDNA was purified using the Wizard DNA Clean-Up Kit (Promega) according to the manufacturer’s instructions.

Purified DNA was submitted to the Biohub-San Francisco for Illumina library preparation and sequencing. Sequencing reads were subsampled to 2 million reads using seqtk v1.3-r106^64^, followed by quality filtering and adapter trimming with fastp v0.20^65^. *De novo* genome assembly was performed using SPAdes v3.13.0^66^. Intergenomic nucleotide similarity between phage genomes was calculated using VIRIDIC^67^ (BLASTn-based) with default parameters (word size = 7, reward = 2, penalty = 3, gap open = 5, gap extend = 2)^68^. Shared gene content was determined using proteinortho v6 with the options --tblastx+ --singles, a minimum percent identity threshold of 25%, and a minimum alignment coverage of 50%^69^. Phage genomes were annotated using Phold^70^. Briefly, predicted open reading frames from *E. lenta* phage contigs were translated and submitted to Phold with default parameters, and the resulting structural homology–based annotations were used to refine phage genome boundaries and assign genes to functional modules.

#### Transmission electron microscopy (TEM)

We performed phage particle TEM following a previously published protocol^71^. Briefly, 1 mL of phage lysate was centrifuged at 25,000 × *g* for 1 h at 4°C. The supernatant was discarded, and the phage pellet was gently resuspended in 0.1 M ammonium acetate and centrifuged again under the same conditions. This washing step was repeated once more, after which the final pellet was resuspended in 0.1 M ammonium acetate. Plaque-forming units (PFUs) were quantified from the cleaned and concentrated phage preparation prior to submission to the University of California, Berkeley Electron Microscopy Facility. Phages were then adsorbed onto transmission electron microscopy grids, fixed with glutaraldehyde, and negatively stained with 0.5% (w/v) uranyl acetate. The resulting grids were inspected using an FEI Tecnai 12 transmission electron microscope (FEI Thermo Fisher, Eindhoven, Hillsboro, OR, USA), and micrographs were acquired with Gatan DigitalMicrograph (Gatan Inc., Pleasanton, CA, USA).

Bacterial cells were fixed in a solution containing 0.1% Ruthenium Red and 2.5% glutaraldehyde/formaldehyde in 0.1 M sodium cacodylate buffer, pH 7.2 for at least 30 min, then stabilized in 1% low-melting-point agarose. The agarose was diced into ∼0.5 mm cubes and fixed overnight. Samples were rinsed (3x, 10 min, RT) in 0.1% Ruthenium Red in 0.1 M sodium cacodylate buffer, pH 7.2, and immersed in 0.1% Ruthenium Red with 1% osmium tetroxide and 1.6% potassium ferricyanide in 0.1 M sodium cacodylate buffer for 1 h. Samples were rinsed again (3×, 10 min, RT) in 0.1% Ruthenium Red in 0.1 M sodium cacodylate buffer, pH 7.2, then subjected to an ascending acetone gradient (10 min each; 35%, 50%, 70%, 80%, 90%, 100%), followed by pure acetone (2×, 10 min, RT). Samples were progressively infiltrated with Epon resin on a rocker and polymerized at 60°C for 24-48 h. Thin sections (90 nm) were cut using a Leica UC6 ultramicrotome (Leica, Wetzlar, Germany) and collected onto formvar-coated copper 50-mesh grids. Grids were post-stained with 2% uranyl acetate followed by Reynolds’ lead citrate for 5 min each. Sections were imaged using a Tecnai 12 120 kV TEM (FEI, Hillsboro, OR, USA), and data were recorded using a Gatan Rio16 CMOS camera with GWS software (Gatan Inc., Pleasanton, CA, USA).

#### Plaque assays

For all plaque assays and phage host range screening experiments, *E. lenta* strains were revived from glycerol stocks in BHI^CHAV^ medium and incubated anaerobically at 37°C for 72 h in a Coy anaerobic chamber. Prior to plating, cultures were vortexed briefly and optical density at 600 nm (OD_600_) was measured to ensure sufficient lawn density (OD₆₀₀ > 0.5). For spot assays, 200 µL of *E. lenta* culture was mixed with 3 mL of sterile 0.2% agarose maintained at 55 °C in a water bath. The mixture was gently inverted to mix and poured onto BHI^CHAV^ agar plates. After solidification of the top agar, 6 µL of serially diluted phage lysate was spotted onto the surface and allowed to air dry. For spread (overlay) assays, 200 µL of *E. lenta* culture was mixed with 200 µL of phage lysate (∼10^9^ PFU/mL) and 3 mL of sterile 0.2% agarose. The mixture was gently mixed and poured onto BHI^CHAV^ agar plates. Plates from both assays were incubated upright at 37°C for 24–48 h under anaerobic conditions before plaque formation was assessed and plaques were enumerated.

All *E. lenta* strains used in phage host range screening are listed in the **Key Resources Table**. Strains that did not produce plaques at the highest phage concentration tested were classified as non-hosts.

#### Gnotobiotic mouse experiments

For the *E. lenta* mono-association experiment, GF C57BL/6J mice were gavaged with 100 µL of *E. lenta* DSM2243 cells resuspended in PBS (10⁸ CFU/mouse; n=9 mice/group). After a 1-week adaptation period, mice were gavaged with 200 µL of either heat-killed ΦKL11 (HK ΦKL11, 0 PFU) or active ΦKL11 (∼10^8^ PFU) resuspended in saline magnesium (SM) buffer. Heat-killed ΦKL11 was prepared by aliquoting lysate into 1.5 mL microcentrifuge tubes and incubating at 90 °C for 3 h in a heat block. Fecal pellets were collected at 24, 48, 72, and 96 h (endpoint) for bacterial and phage load quantification. Briefly, fecal pellets were resuspended in PBS to a concentration of 100 mg/mL. For bacterial load quantification, fecal suspensions were serially diluted in PBS and spread-plated onto BHI^A^ agar (100 µL per plate per dilution). For phage load quantification, spread plaque assays were performed as described above. Stomach and cecum contents were collected in addition to fecal pellets at 96 h post-gavage for bacterial and phage load quantification. *E. lenta* abundance was determined as colony-forming units (CFU) per gram stool. ΦKL11 abundance in fecal pellets, stomach, and cecum was determined as plaque-forming units (PFU) per gram contents.

We also tested the impact of ΦKL11 on *E. lenta* in an independent mouse genetic background. GF male BALB/c mice were gavaged with 200 µL of *E. lenta* DSM2243 resuspended in PBS (∼10⁹ CFU per mouse; n = 3–4 mice per group), while GF control mice received 200 µL of PBS. After a 1-week adaptation period, mice were gavaged with 200 µL of heat-killed ΦKL11 (HK ΦKL11; no detectable PFU) or active ΦKL11 (∼10¹⁰ PFU), both resuspended in saline magnesium (SM) buffer. Heat-killed ΦKL11 was prepared as previously described. Fecal pellets were collected daily from day 0 to day 6 for quantification of bacterial and phage loads as previously described.

For the *E. lenta* DSM2243 and *E. lenta* DSM15644 co-colonization experiment, GF BALB/c male mice were gavaged with a 50:50 mixture of both strains (∼10⁸ CFU each). The heat-killed (HK) ΦKL11 group (n = 10) and the ΦKL11 group (n = 9) received either heat-inactivated phage (no detectable PFU) or active phage (10⁸–10⁹ PFU/mL) in drinking water, starting three days after *E. lenta* administration and continuing for two consecutive weeks. Fecal samples were collected at 1, 2, 3, 4, 7, 9, 11, and 14 days after phage dosing (day 0).

For the synthetic community experiment, a four-member community—*E. lenta* DSM2243, *B. thetaiotaomicron* VPI-5482, *E. coli* MG1655, and *C. innocuum* DSM 1286—was mixed at a 1:1:1:1 ratio and administered by oral gavage to BALB/c GF mice at 10⁸ CFU/mL (200 µL per mouse). After a 1-week acclimation, mice received either 0 PFU/mL heat-killed (HK) or 10^8^ PFU/mL active ΦKL11 via drinking water (n=5 mice/group). Mice were euthanized on day 13. Stool samples were collected on 0, 1, 2, 3, 6, 7, 8, 9, 10, and 13 after phage dosing (day 0). For both co-culture and synthetic community experiments, *E. lenta* abundance was determined by qPCR on gDNA extracted from mouse fecal samples and expressed as genome equivalents per gram of feces.

#### gDNA extraction from mouse fecal samples

We weighed each fecal pellet and used 10 to 60 mg of each sample for DNA extraction using the ZymoBIOMICS 96 MagBead DNA Kit (D4302). Briefly, 650 μl of the lysis buffer was added to the sample, then incubated at 65 °C for 10 minutes prior to 5 min bead beating using Biospec Mini-Beadbeater^TM^. After this, 200 μl of the mixture was added to a 2 mL 96-well deep-well block and 600 μl of binding buffer and 25 μl of magnetic beads were added. This was vortexed at 750 rpm for 10 minutes, the beads were captured on a magnetic stand, and the supernatant discarded. The beads were washed with 900 μl Wash 1 buffer once and twice with 900 μl Wash 2 buffer. After aspiration of the supernatant, the plate was spun briefly and dried at 65 °C for ∼30 minutes. The DNA was eluted with 100 μl of water and DNA kept at −20 °C.

#### Probe-based qPCR for strain abundance quantification

Bacterial gDNA was diluted 1:10 in nuclease-free water. Reactions were performed in 10 µL total volume containing 4 µL diluted DNA and 6 µL master mix (SsoAdvanced Universal Probe Supermix) supplemented with oligonucleotides and probes according to the manufacturer’s instructions. For bacterial quantification in co-colonization experiments, bacterial abundance was measured by fluorophore-based quantitative PCR. Total bacterial load was determined using a universal primer set that amplifies a conserved bacterial target present in both strains. The abundance of *E. lenta* DSM2243 was quantified in parallel using a DSM2243-specific primer set targeting a strain-specific locus. Quantification of the four bacterial species was performed using a combination of singleplex and triplex probe-based qPCR assays (oligonucleotide and probe sequences in **Table S6**). qPCR was run on a Bio-Rad CFX384^TM^ Real-Time System using the following cycling conditions: 95 °C 3 min; 40 cycles of 95 °C 10 s and 60 °C 30 s. Samples were run in technical triplicate, and Cq values were averaged prior to analysis. *E. lenta* genome equivalents were determined by standard curve–based absolute quantification using serial dilutions of *E. lenta* gDNA. Cq values were interpolated onto the standard curve and converted to genome equivalents, with values corrected for dilution and normalized to sample mass.

#### Recovery of *E. lenta* from distal gut samples

1-2 fecal pellets per mouse from the *E. lenta* C57BL/6J mono-association experiment were resuspended in PBS and subcultured into BHI^A^ medium (n=4 mice/group). After 72 h of incubation in an anaerobic chamber, cultures were converted to glycerol stocks by mixing 300 µL of sterile 50% glycerol with 700 µL of culture in a cryogenic vial. These stocks were designated passage 1 (P1). All *in vitro* analyses of mouse-recovered *E. lenta* were derived from this P1 stock. To passage the *E. lenta* recovered from feces, an aliquot of P1 was inoculated into fresh BHI^A^ and incubated anaerobically for 72 h, then subcultured; this 72 h growth/subculture cycle was repeated to generate passages P2-P4.

#### *In vitro* phage selection

*E. lenta* cells (∼10^6^ CFU) were mixed with ΦKL11 (∼10 PFU) in BHI^A^ to generate passage 1 (P1). After 48-72 h of incubation in an anaerobic chamber, 200 µL of culture was plated onto BHI^A^ agar to obtain passage 2 (P2). Colonies from P2 were subjected to two additional rounds of co-culture with ΦKL11 in BHI^A^ to generate P3 and P4. For P1, P3, and P4, susceptibility to ΦKL11 was assessed by plaque (spot) assay, and phage abundance in culture supernatants was quantified. To prepare overlays, *E. lenta* DSM2243 wild-type culture was mixed with 0.2% agarose. Supernatants from P1, P3, and P4 were filtered through 0.22 µm filters, serially diluted in SM buffer, and spotted onto the bacterial overlays. Plaques were counted after 48 h of incubation at 37 °C in an anaerobic chamber.

#### Single-cell sorting

*E. lenta* DSM2243 wild-type was subcultured from a single colony into BHI^A^ medium and grown for 36-48 h. Cultures were harvested at an OD₆₀₀ of 0.4-0.6. Cells were washed once with PBS and diluted to an OD₆₀₀ of 0.05 in PBS. To prepare the cell-recovery plate, 200 µL of pre-equilibrated BHI^A^ supplemented with 10 mM sodium formate was dispensed into each well of a 96-well plate inside an anaerobic chamber and sealed with heat-sealing foil. Single-cell sorting was performed using the Cytena B.SIGHT™ on the benchtop under aerobic conditions. The gating parameters were set to a size range of 0.5-4 µm and a roundness range of 0-0.5. After sorting, the plate was returned to the anaerobic chamber. Two hundred microliters of each single-cell-dispensed culture was transferred into a deep 96-well plate containing 800 µL of BHI^A^ supplemented with 10 mM sodium formate. Plates were sealed with a Breathe-Easy sealing membrane to facilitate gas exchange and incubated for 72 h at 37 °C in the anaerobic chamber.

#### Bacterial gDNA extraction, sequencing, and assembly

For genomes obtained from Pacbio sequencing (**Table S4**), 40 mL of BHI^CHAV^ broth was inoculated with a culture of *E. lenta* and incubated for 72 h under standard conditions. Firstly, the bacterial culture underwent centrifugation with the resulting bacterial pellet resuspended in NucleoBond Buffer G3 of NucleoBond Buffer Set III (Macherey-Nagel, Düren, Germany), supplemented with RNase, lysozyme and proteinase K, and incubated at 37 °C for 16-18 hrs. Buffer G4 of NucleoBond Buffer Set III of the same kit was added and incubated at 50 °C for 1 hrs and centrifuged to remove insoluble material. High molecular weight gDNA was isolated from lysate utilising Nucleobond AXG100 (Macherey-Nagel) columns in accordance with the manufacturer’s protocol. Preparation of gDNA for PacBio sequencing utilized the HiFi SMRTbell library kit and a BluePippinsize selection (Beverly, MA, USA) on the pool of the libraries. Sequencing was performed with 8M SMRT cells on Sequel II (Pacific Biosciences). Genome assembly of circular genomes was conducted with Flye (v2.9-b1768), with Minimap2 (v2.17) and Samtools (v1.7) used for the calculation of sequence coverage^72–74^.

For the genome obtained using MinION (Oxford Nanopore Technologies) (**Table S4**), gDNA was isolated using the GenElute™ Bacterial gDNA Kit. DNA was then quantified using the Qubit broad range assay before standardization for paired-end Nextera XT library preparation. Libraries were inspected with the TapeStation 4200 system (Agilent, Santa Clara, CA, USA) using the High Sensitivity D1000 ScreenTape Assay before sequencing with an Illumina NovaSeq platform (Illumina Inc., San Diego, CA, USA). Long-read gDNA libraries were prepared with an Oxford Nanopore SQK-LSK109 kit with Native Barcoding EXP-NBD104 (Oxford Nanopore Technologies, Oxford, UK). Barcoded samples were pooled together into a single sequencing library and loaded in an FLO-MIN111 (R.10.3) flow cell in a MinION (Oxford Nanopore Technologies). Genome assembly of circular genomes was conducted with Unicycler (v0.4.8) using short and long reads obtained from Illumina and Nanopore sequencing, respectively^75^. Short and long read sequence coverage was determined with Bowtie2 (v2.3.4.1) and minimap2 (v2.17), respectively, in conjunction with Samtools (v1.7)^76^.

For genomes obtained from the germ-free monoassociation experiment (**Figs. 1D–G**), stool-derived *E. lenta* were grown anaerobically in liquid culture for 2–3 days to OD₆₀₀ ≈ 0.7–0.8. For each sample, cells from 10 mL of culture were pelleted, washed once with PBS, and resuspended in 1.5 mL lysis solution from the ZymoBIOMICS 96 MagBead DNA Kit (D4308). Cell suspensions were split across two bead-beating columns and mechanically lysed at 2000 rpm for 5 min, followed by 5 min incubation on ice. Lysates were clarified by centrifugation at 10,000 × *g* for 1 min, and the supernatant was transferred to a new tube and further centrifuged at 4,000 × *g* for 5 min to remove residual debris. Clarified lysates (∼400–600 µL per sample) were stored at −80°C until DNA extraction. For gDNA purification, 200 µL of lysate was processed per microcentrifuge tube following the manufacturer’s instructions. For each sample, two parallel extractions were performed, and DNA was eluted in a combined total volume of 50 µL nuclease-free water. Purified gDNA was shipped on dry ice to SeqCenter (Pittsburgh, PA, USA) for Oxford Nanopore MinION sequencing.

#### Identification of structural variants

Structural variants (SVs) in *Eggerthella lenta* strains were identified from long read sequencing data generated from pure cultures using either Oxford Nanopore or PacBio platforms. Long reads were aligned to either the primary genome assembly generated from the same dataset or to a reference genome using MiniMap2 (v2.17)^72,77^. SVs were detected using Sniffles (v2.0.7)^77^ based on split read and discordant alignment signatures relative to the reference assembly. Reads spanning predicted breakpoint regions were extracted using Samtools (v1.7) and SeqKit (v2.2.0)^73,78^. Structural variant calls and supporting reads were manually inspected in Artemis (v16) at the genomic coordinates reported by Sniffles to confirm inversion boundaries and read orientation^79^. To reconstruct alternative genomic configurations, only reads supporting the variant structure were selected and assembled de novo using Flye (v2.9-b1768)^74^. The resulting assemblies were compared with the primary assembly to confirm the structure and boundaries of the inversion events.

#### Quantification of SV abundance by qPCR

gDNA used for qPCR was prepared from *E. lenta* cultures as described above and normalized to 5 ng/µl. The relative abundance of each structural variant (SV) in *E. lenta* DSM2243 was quantified using SV-specific primer sets (**Table S6**) with PowerUp™ SYBR™ Green Master Mix on a Bio-Rad CFX384™ Real-Time PCR System. Primer specificity and amplification efficiency were initially evaluated using amplicons cloned into TOPO vectors. TOPO constructs were generated using the TOPO TA Cloning Kit and transformed into Mach1™-T1ᴿ One-Shot™ Cells according to the manufacturer’s instructions. qPCR was performed with the following thermocycling conditions: UDG activation at 50 °C for 2 min, initial denaturation at 95 °C for 2 min, followed by 35 cycles of 95 °C for 15 s, 65 °C for 15 s, and 72 °C for 15 s. SV abundance was determined using a relative quantification approach normalized to a single-copy housekeeping locus, *elnmrk*^40^. For each sample, ΔCq was calculated as Cq(SV) − Cq(*elnmrk*), and relative SV abundance was expressed as 2^−ΔCq^.

#### SV allele frequency analysis

Long-read sequences were aligned to the reference genome using Minimap2 v2.17 (pre-set ONT parameters) and indexed using SAMtools v.1.7 (default parameters). The read alignments were analyzed using Sniffles2 (with minimum support of 5 and minimum length of 500 bp) to determine SV frequencies.

#### Genome alignment visualization

Whole-genome alignments were visualized using EasyFig (v2.2.5) on a Windows platform. GenBank files were used as input for alignment. Pairwise sequence similarity was computed using BLASTn within EasyFig with the following parameters: minimum alignment length of 2,000 bp, maximum E-value of 1 × 10⁻⁵, and minimum percent identity of 70% for *E. lenta* DSM2243 and minimum alignment length of 100 bp, maximum E-value of 1 × 10⁻^4^, and minimum percent identity of 70% for *E. lenta* 539-5C, 943-4, and FCC8. Alignment outputs were displayed to illustrate genome structure conservation and rearrangements across strains.

#### Bacterial RNA extraction

All cultures were grown anaerobically in Hungate tubes containing liquid BHI^A^. Stationary-phase cultures were split into pairs and diluted to an OD₆₀₀ of 0.08–0.10 in a total culture volume of 3 mL. Cultures were then grown overnight and monitored until they reached an OD₆₀₀ of 0.3. Total RNA was extracted using the Qiagen RNeasy Kit, with minor modifications optimized for our strains. Briefly, 1.5 mL of culture at an OD₆₀₀ of 0.3 was collected and mixed with 3 mL of RNAprotect Bacteria Reagent. Samples were centrifuged at >16,000 × *g* for 10 min and the supernatant was carefully removed by pipetting. Pellets were flash-frozen in liquid nitrogen and stored at −80 °C for extraction the following day. For lysis, pellets were thawed on ice and resuspended in 900 µL Buffer RLT supplemented with 1% β-mercaptoethanol, transferred to 2 mL MP Lysing Matrix tubes, and lysed for 50 s in a Biospec Mini-Beadbeater 96. Tubes were centrifuged at maximum speed (∼21,000 × *g*) for 30 s, and 700 µL of each clarified lysate was transferred to new microcentrifuge tubes on ice. An equal volume (700 µL) of 70% ethanol was added to each lysate and mixed by pipetting. Then, 700 µL of the lysate/ethanol mixture was loaded onto an RNeasy spin column, centrifuged for 30 s at 10,000 × *g*, and the flow-through discarded; this step was repeated with the remaining 700 µL of lysate. The remainder of the protocol was performed according to the manufacturer’s instructions, including on-column DNase digestion, except that RNA was eluted in 100 µL RNase-free water.

#### RNAseq and data processing

Samples were treated with Invitrogen DNase I (RNase-free) according to the manufacturer’s instructions. Libraries were prepared using the Illumina Stranded Total RNA Prep Ligation with Ribo-Zero Plus kit and indexed using 10-bp dual indices manufactured by Integrated DNA Technologies (IDT) for Illumina. Sequencing was carried out on a NextSeq 2000, generating 2 × 50 bp paired-end reads. Demultiplexing, quality control, and adapter trimming were performed by SeqCenter (Pittsburgh, PA) using Illumina bcl2fastq (v2.20.0.445) with default parameters. RNA-seq reads were mapped to the *E. lenta* reference genome using Bowtie2 (v2.3.4.1). Alignment (SAM) files were processed with Samtools (v1.7) to generate BAM files, which were then used to create gene-level count matrices normalized by gene length. Differentially expressed gene (DEG) analysis was performed in R using the DESeq2 (v1.44.0) package as part of the Bioconductor distribution. As a preprocessing step, genes with zero counts across all samples were removed from the count matrices. DEG analysis was then conducted using the DESeq function in DESeq2.

#### Comparative genomic analyses of ΦKL11 susceptibility

Orthologous gene content was associated with ΦKL11 susceptibility using a presence–absence–based genome-wide analysis across 102 *Eggerthella lenta* genomes (**Table S11**). Orthologous groups were identified using eggNOG-mapper v2 and collapsed into a binary gene presence/absence matrix across strains. ΦKL11 susceptibility was encoded as a binary phenotype (host or non-host). For each orthologous group, association with susceptibility was tested using Fisher’s exact test on 2 × 2 contingency tables, excluding groups lacking variation in gene presence or phenotype. Effect sizes were summarized as the difference in gene prevalence between sensitive and resistant strains. Genes with a nominal p-value < 0.01 were considered candidates for association.

Separately, we performed a targeted analysis of capsule polysaccharide (CPS) loci across the same 102 *E. lenta* genomes (**Table S11**). eggNOG ortholog annotations were used to visualize the distribution of CPS-associated genes across strains (**Fig. S10A**). Each CPS cluster was considered present if at least 85% of CPS cluster genes (as called by eggNOG orthology) were present across the reference genome, regardless of genomic location. Associations between CPS cluster presence and ΦKL11 susceptibility were tested using Fisher’s exact test (*p* < 0.05).

#### Phylogenetic tree construction

A phylogenetic tree of *Eggerthella lenta* genomes was constructed using PhyloPhlAn v3.1.194^80^ based on conserved marker genes. Phylogenomic analysis was performed using the *E. lenta*-specific marker database with amino acid sequence alignments under the low-diversity parameter setting. The resulting alignment was used to infer a maximum-likelihood phylogeny. *Eggerthella timonensis* Marseille-P3135 was included as an outgroup to root the tree.

## Acknowledgments

We thank the UCSF Gnotobiotics Core for assistance with mouse experiments and Dr. Danielle Jorgens and Reena Zalpuri at the University of California Berkeley Electron Microscope Laboratory for advice and assistance in electron microscopy. Sequencing was performed at Biohub, San Francisco, the UCSF Center for Advanced Technology, and the Norwegian Sequencing Centre, University of Oslo, Norway. Funding was provided by the National Institutes of Health (R01AT011117, R01HL122593, R01CA255116, and R01DK114034, P.J.T.; F32GM140808, C.N.), the Canadian Institutes of Health Research (K.N.L.), and the Government of Ireland Postdoctoral Fellowship 2019 (project ID GOIPD/2019/1097, C.B.), and European Research Council (ERC), under the European Union’s Horizon 2020 research and innovation programme (grant agreement No. 101001684 – PHAGENET, A.S.). P.J.T is a Biohub, San Francisco, Investigator and held an Investigators in the Pathogenesis of Infectious Disease Award from the Burroughs Wellcome Fund. Diagrams were created with Biorender.com.

## Author contributions

S.Z. and C.B. contributed to conceptualization, data curation, formal analysis, investigation, methodology, and writing of the original draft, and participated in writing – review & editing. K.R.T., K.N.L., L.R.H., P.S.-P., and P.C. contributed to formal analysis, investigation, and methodology; K.R.T. additionally contributed software. E.O., J.L., L.R., G.P., and T.L. contributed to investigation. C.N. contributed to methodology. F.B. contributed to methodology, software, and writing – review & editing. A.S., L.A.D., and P.R. contributed to supervision; A.S. and P.R. also contributed to methodology and writing – review & editing, and L.A.D. contributed to writing – review & editing. A.C. contributed to funding acquisition and writing – review & editing. C.H. and P.J.T. contributed to conceptualization, funding acquisition, methodology, supervision, and writing – review & editing.

## Declaration of interests

P.J.T. is on the scientific advisory boards of Pendulum and SNIPRbiome. All other authors declare no competing interests.

## Declaration of generative AI and AI-assisted technologies

During the preparation of this manuscript, the authors used ChatGPT5.2 (OpenAI) to assist with editing for improved clarity and readability. After using this tool, the authors extensively revised the content and they accept full responsibility for the content of the publication.

## SUPPLEMENTAL FIGURES AND LEGENDS

**Figure S1.**
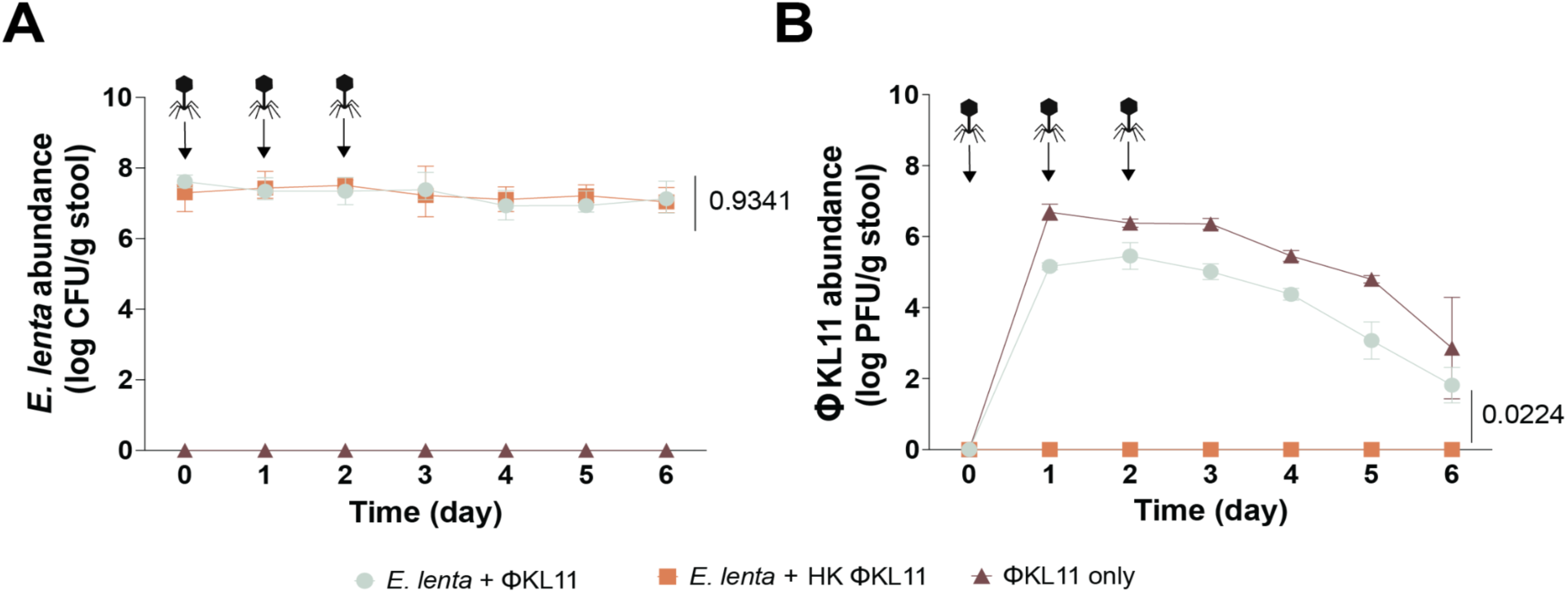
*E. lenta* evades phage predation in the mouse gut in a repeated experiment. **(A–B)** Germ-free (GF) BALB/c mice were monoassociated with *E. lenta* DSM2243 for one week prior to phage treatment. Mice then received either heat-killed (HK) or active ΦKL11 by oral gavage for three consecutive days. A GF control group that received active ΦKL11 only was also included (n = 3-4 mice per group). **(A)** Longitudinal stool measurements of *E. lenta* abundance (CFU/g) and **(B)** ΦKL11 abundance (PFU/g). *p*-values, two-way ANOVA.

**Figure S2.**
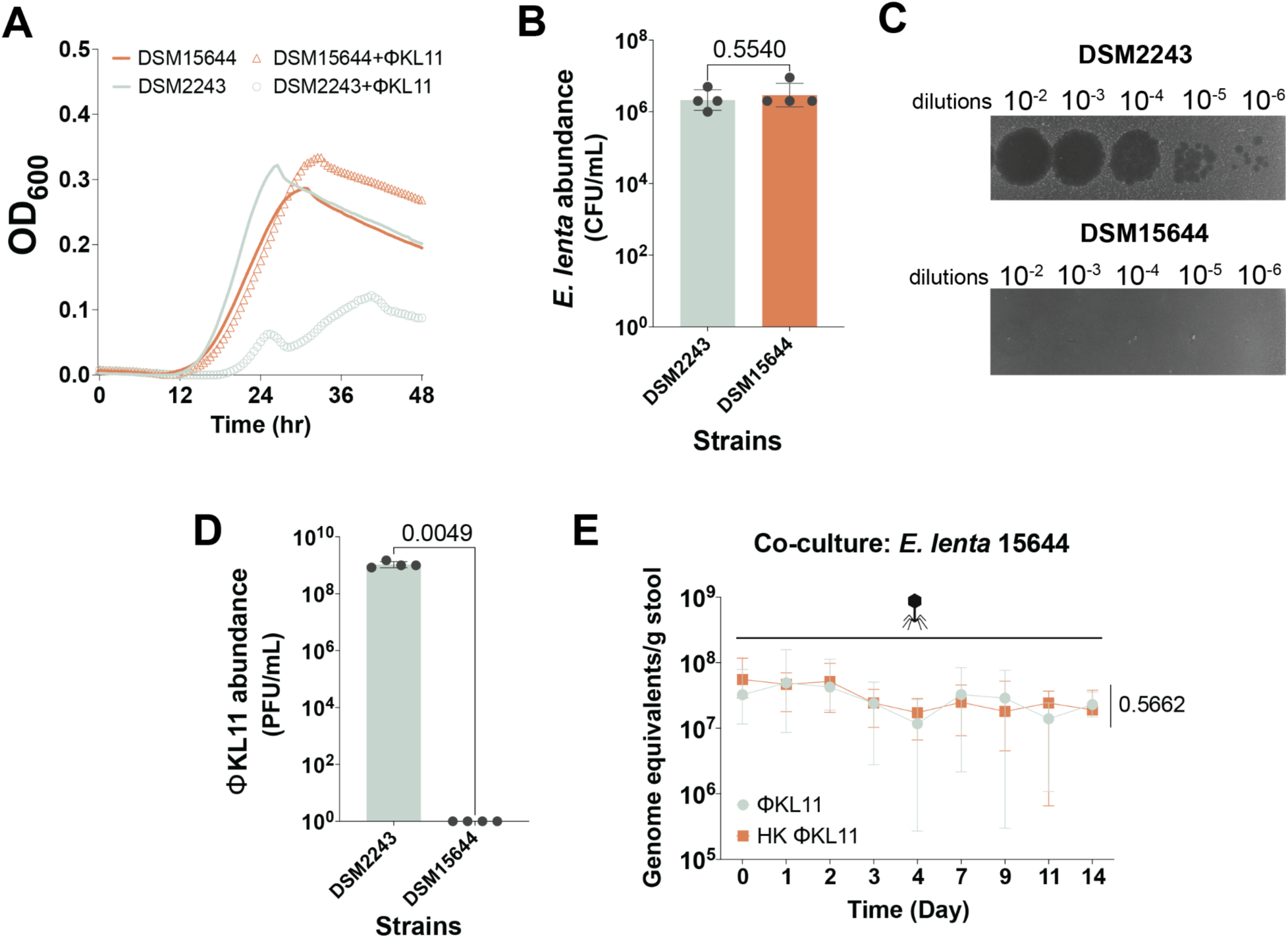
*E. lenta* evades phage predation in a co-culture experiment in GF mice. **(A)** Growth curves of *E. lenta* DSM2243 and *E. lenta* DSM15644 in the presence or absence of ΦKL11. Curves represent the mean of *n*=4 biological replicates. **(B)** *E. lenta* cell counts for DSM2243 and DSM15644 cultures used in the plaque assay in **Fig. S2C**. **(C-D)** ΦKL11 robustly lyses *E. lenta* DSM2243 on an agarose overlay but does not lyse *E. lenta* DSM15644. **(E)** *E. lenta* DSM15644 abundance measured by qPCR during co-colonization with the ΦKL11-sensitive *E. lenta* DSM2243 (1:1 inoculation; n=9–10 mice per group). *p*-values, Welch’s *t* test **(B,D)** and two-way ANOVA **(E)**.

**Figure S3.**
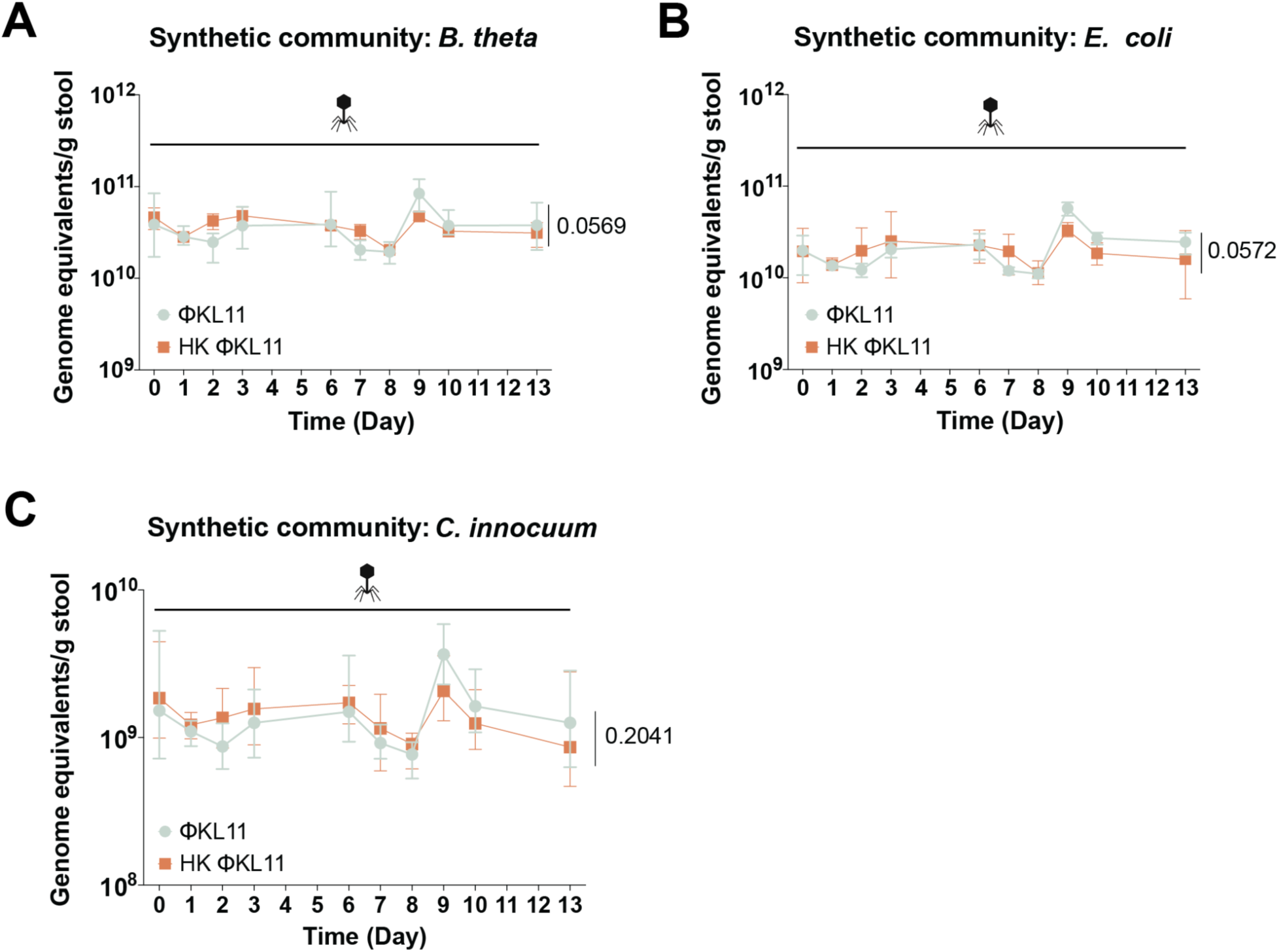
*E. lenta* evades phage predation in a 4-member synthetic microbiota. **(A-C)** Abundance of **(A)** *B. thetaiotaomicron*, **(B)** *E. coli*, and **(C)** *C. innocuum* measured by qPCR during co-colonization with *E. lenta* DSM2243 (1:1:1:1 initial inoculum; n=5 mice/group). *p-*values, two-way ANOVA.

**Figure S4.**
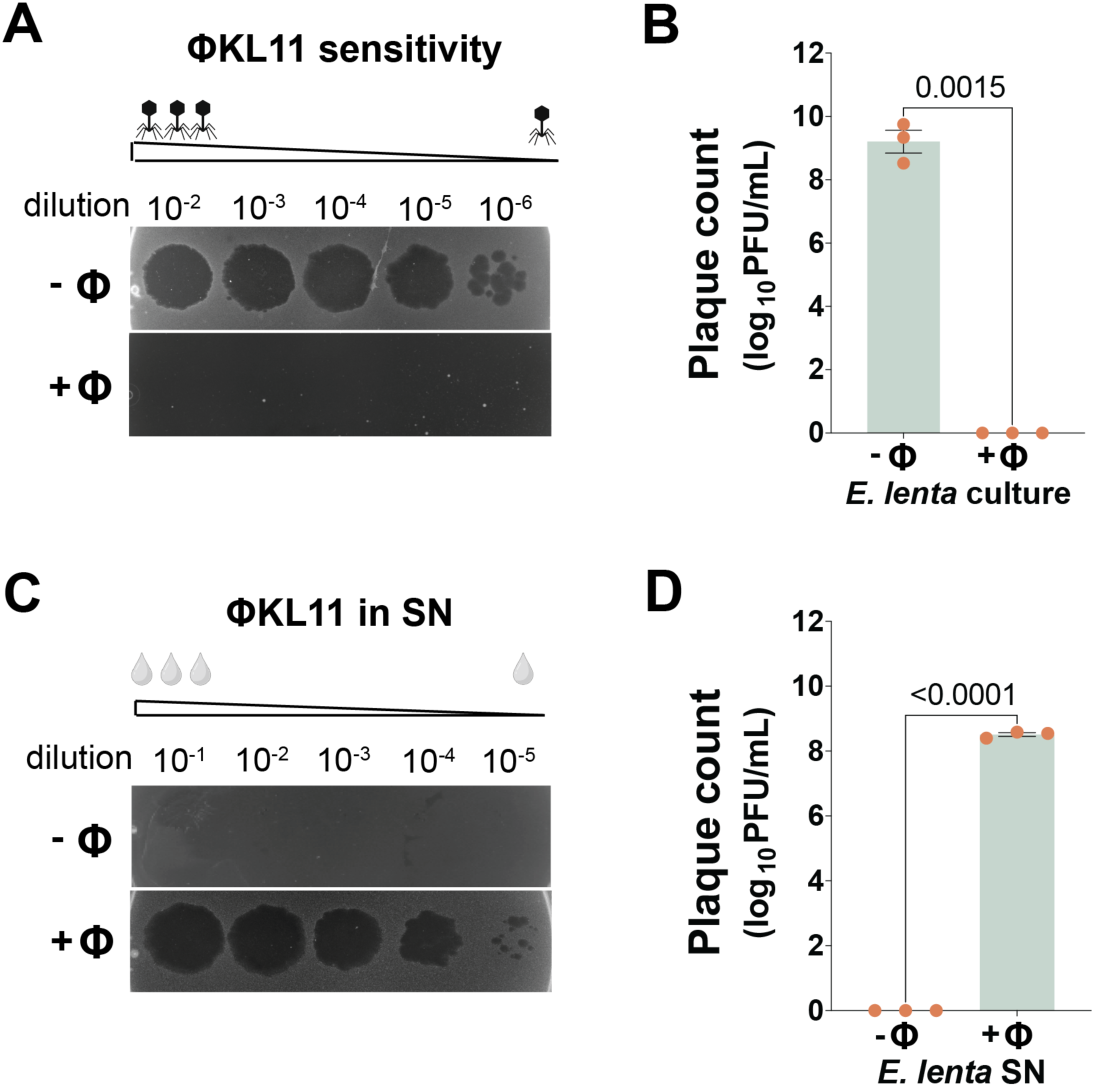
*In vitro* selection for *E. lenta* phage resistance. **(A,B)***E. lenta* DSM2243 develops resistance to ΦKL11 after four serial passages in BHI^A^ liquid broth in the presence of ΦKL11. **(C,D)** ΦKL11 was detected at high levels (8.51 ± 0.10 log₁₀ units) in the supernatant of ΦKL11-exposed *E. lenta* cultures. Values are mean±SEM. *p*-values, Welch’s *t* tests.

**Figure S5.**
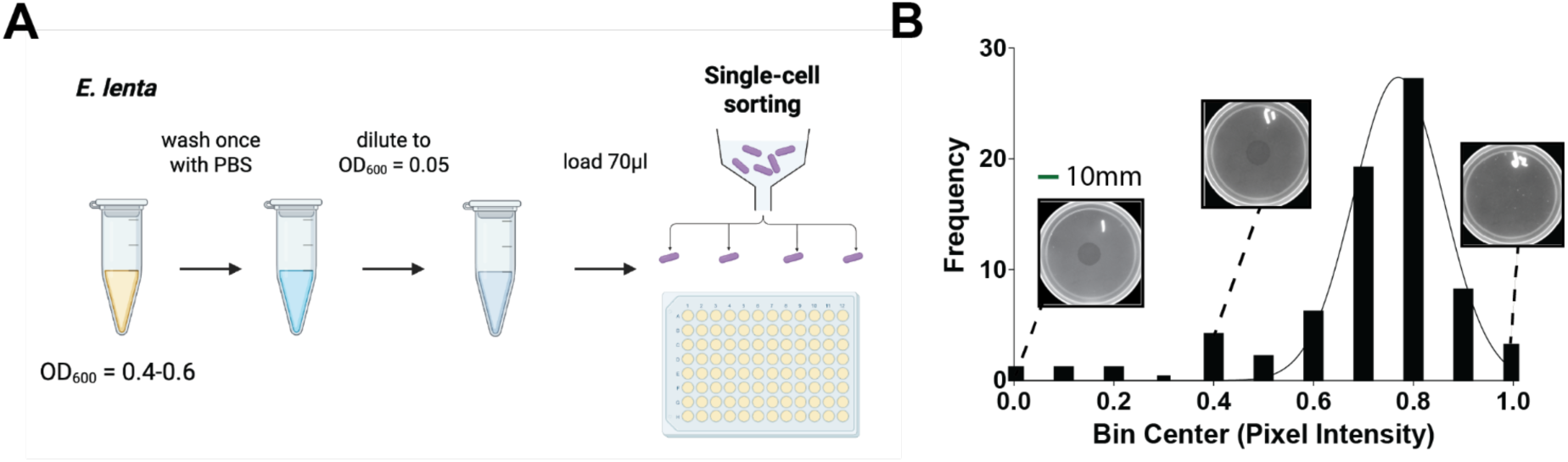
Single cell workflow and distribution of plaque intensities. **(A)** Workflow diagram. **(B)** Histogram showing the distribution of plaque pixel intensities measured from Φ^R1^-derived plaques formed on *E. lenta* DSM2243. The x-axis indicates the bin center of normalized pixel intensity, and the y-axis indicates the frequency of plaques within each bin. Representative plaque images corresponding to low, intermediate, and high intensity bins are shown above the histogram. Scale bar, 10mm.

**Figure S6.**
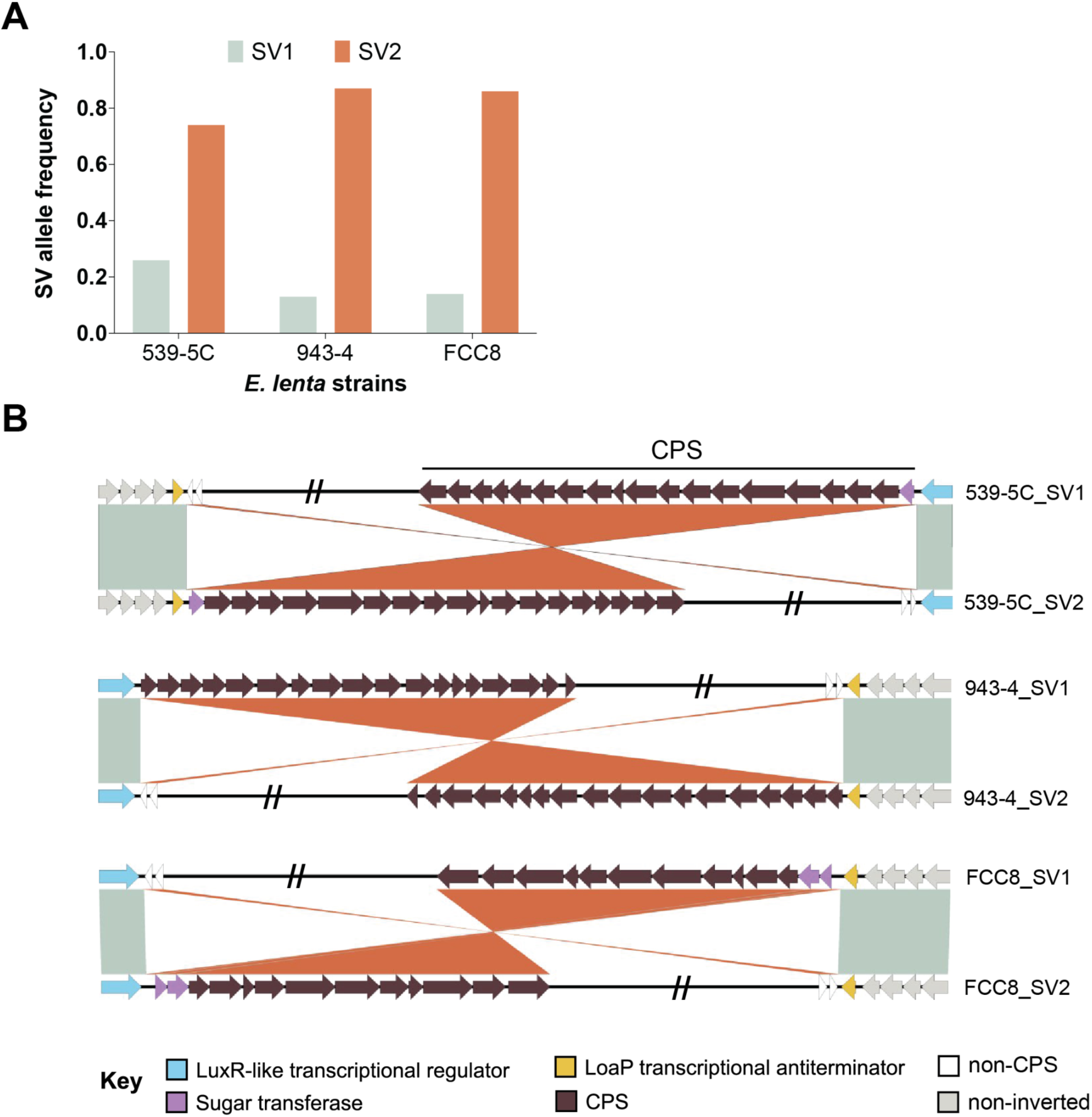
Large genomic inversions are detected in other *E. lenta* strains. **(A)** Relative abundance of SVs in *E. lenta* strains 539-5C, 943-4, and FCC8, as quantified by long-read sequencing. **(B)** Comparative alignment of the flanking genomic neighborhoods of the inversions identified in *E. lenta* 539-5C SV1/2, 943-4 SV1/2, and FCC8 SV1/2. Orange ribbons denote homologous blocks (crossed ribbons indicate inversions), and mint shading marks syntenic regions in the same orientation. Arrowed boxes represent CDS, highlighting key features of each CPS gene cluster (LuxR-like regulator, sugar transferases, LoaP). Double slashes (“//”) denote a 1.9 to 2-Mbp region omitted in SVs.

**Figure S7.**
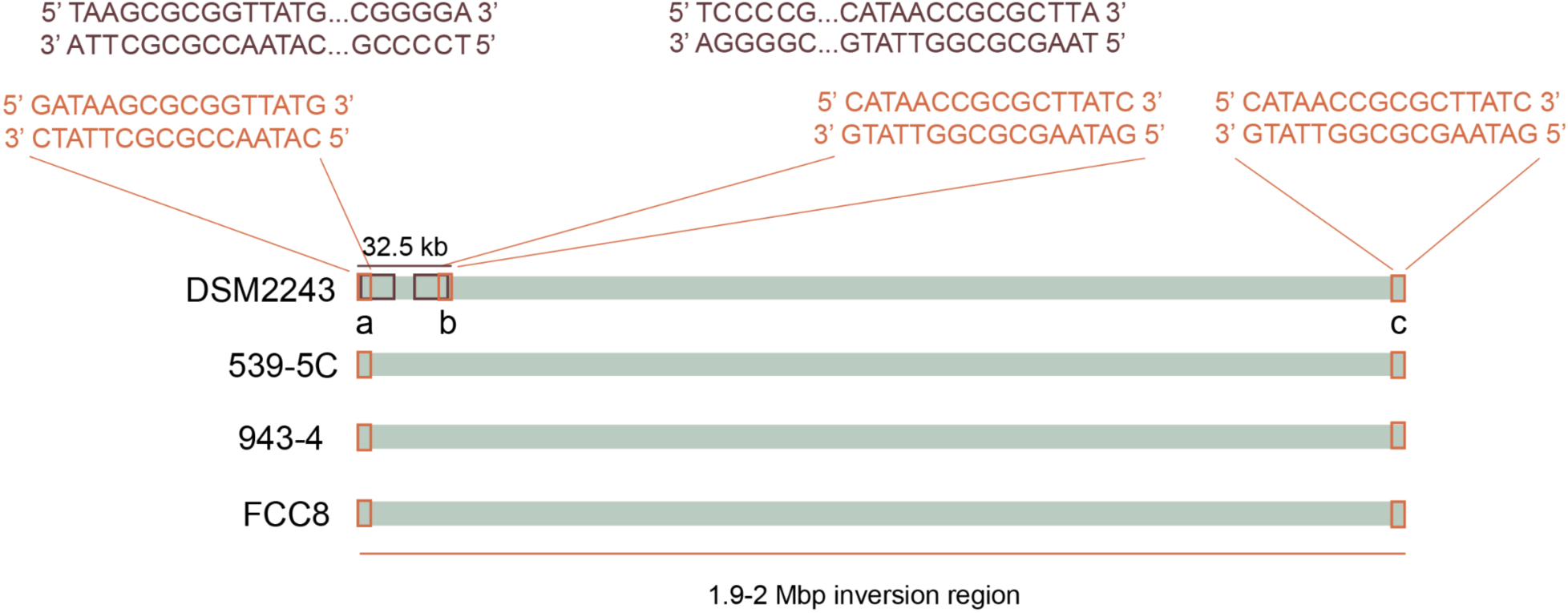
The same 16-bp inverted repeat sequence was identified in *E. lenta* strains harboring large chromosomal inversions. A conserved 16-bp inverted repeat (IR) sequence is observed at inversion junctions a, b, and c in DSM2243 and flanks the ∼2-Mbp inversion in strains 539-5C, 943-4, and FCC8. A 39-bp IR sequence is also detected at inversion junctions a and b in DSM2243, flanking the ∼32.5 kb inversion. The 16-bp IR overlaps the 39-bp IR and extends 2 bp beyond its boundaries. Orange boxes indicate the 16-bp IR sequences, while brown boxes indicate the 39-bp IR sequences.

**Figure S8.**
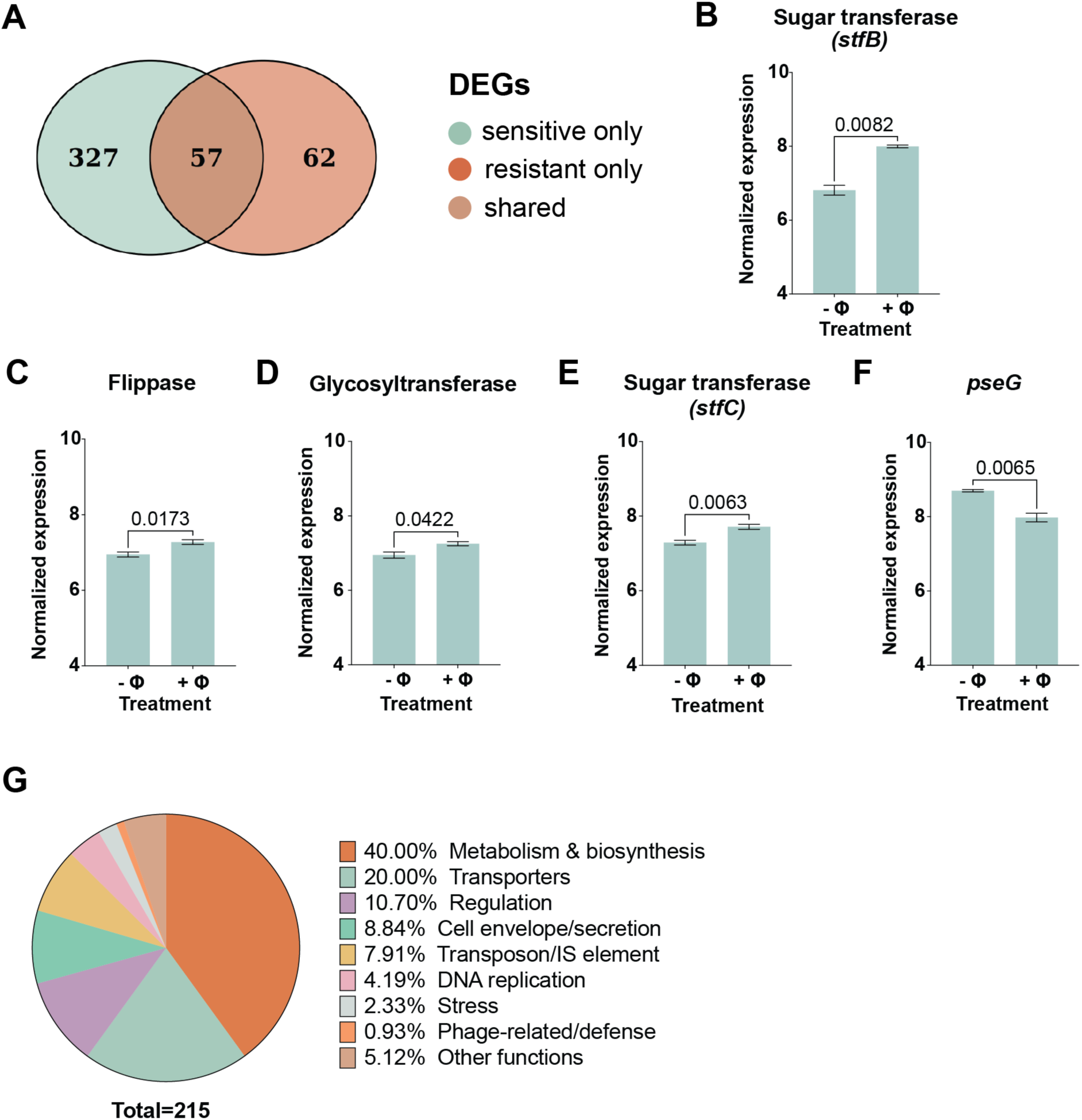
Phage-induced transcriptional responses differ between sensitive and resistant *E. lenta* backgrounds. **(A)** Venn diagram showing the overlap of differentially expressed genes (DEGs) in response to phage exposure in ΦKL11^S^ and ΦKL11^R^ *E. lenta* DSM2243. DEGs were identified by comparing ΦKL11-treated (+Φ) versus no-phage control (-Φ) conditions separately within each host background (*p_adj_*<0.05 and |log₂ fold change|≥1). Numbers indicate genes uniquely responsive to phage in sensitive hosts (left), uniquely responsive in resistant hosts (right), or shared between both backgrounds (overlap). **(B-E)** Genes within the CPS2 cluster that are differentially expressed exclusively in phage-sensitive *E. lenta* upon ΦKL11 exposure, shown as DESeq2 variance-stabilized transformed (VST) expression value: **(B)** sugar transferase *stfB* (ELEN_RS12115), **(C)** flippase (ELEN_RS12140), **(D)** glycosyltransferase (ELEN_RS12240), and **(E)** CPS3 sugar transferase *stfC* (ELEN_RS12255). **(F)** *pseG* (ELEN_RS12180) is significantly down-regulated in ΦKL11-treated ΦKL11^S^ hosts. **(G)** Functional classification of genes up-regulated in ΦKL11^S^ *E. lenta* following ΦKL11 exposure (total genes=215). Differentially expressed genes in response to phage infection were assigned to functional categories based on predicted annotations, and the pie chart indicates the relative distribution of each category. Genes lacking functional annotations (e.g., hypothetical proteins) were excluded from this analysis. Values are mean±SEM; *p*-values, Welch’s *t* tests **(B-F)**.

**Figure S9.**
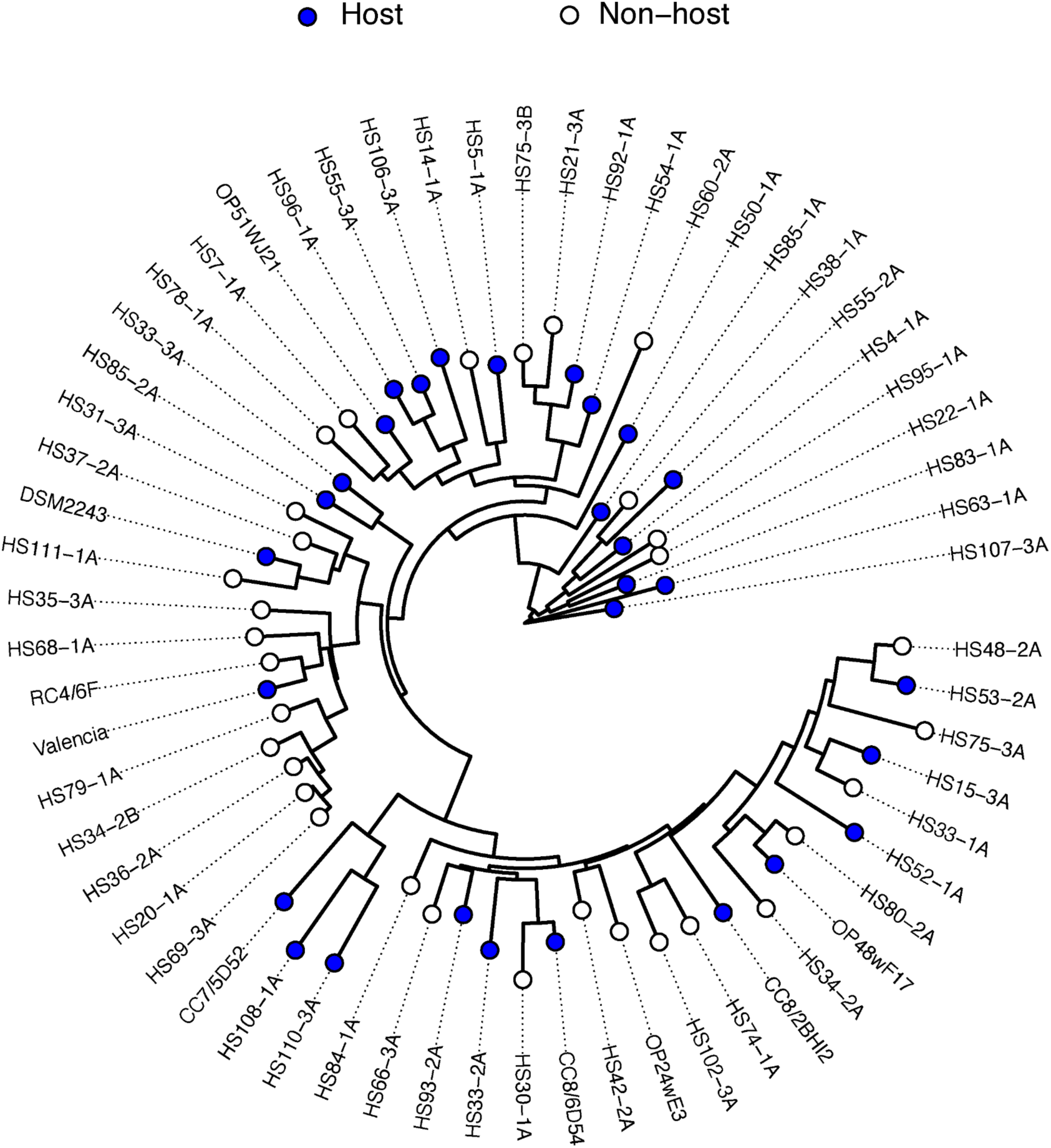
Bacteriophage ΦKL11 has a broad host range. Phylogenetic tree of *E. lenta* strains constructed from core genome alignment. ΦKL11 sensitivity, quantified by plaque assay, is overlaid as a colored dot at each tip, indicating host versus non-host status (n = 61 genomes).

**Fig. S10.**
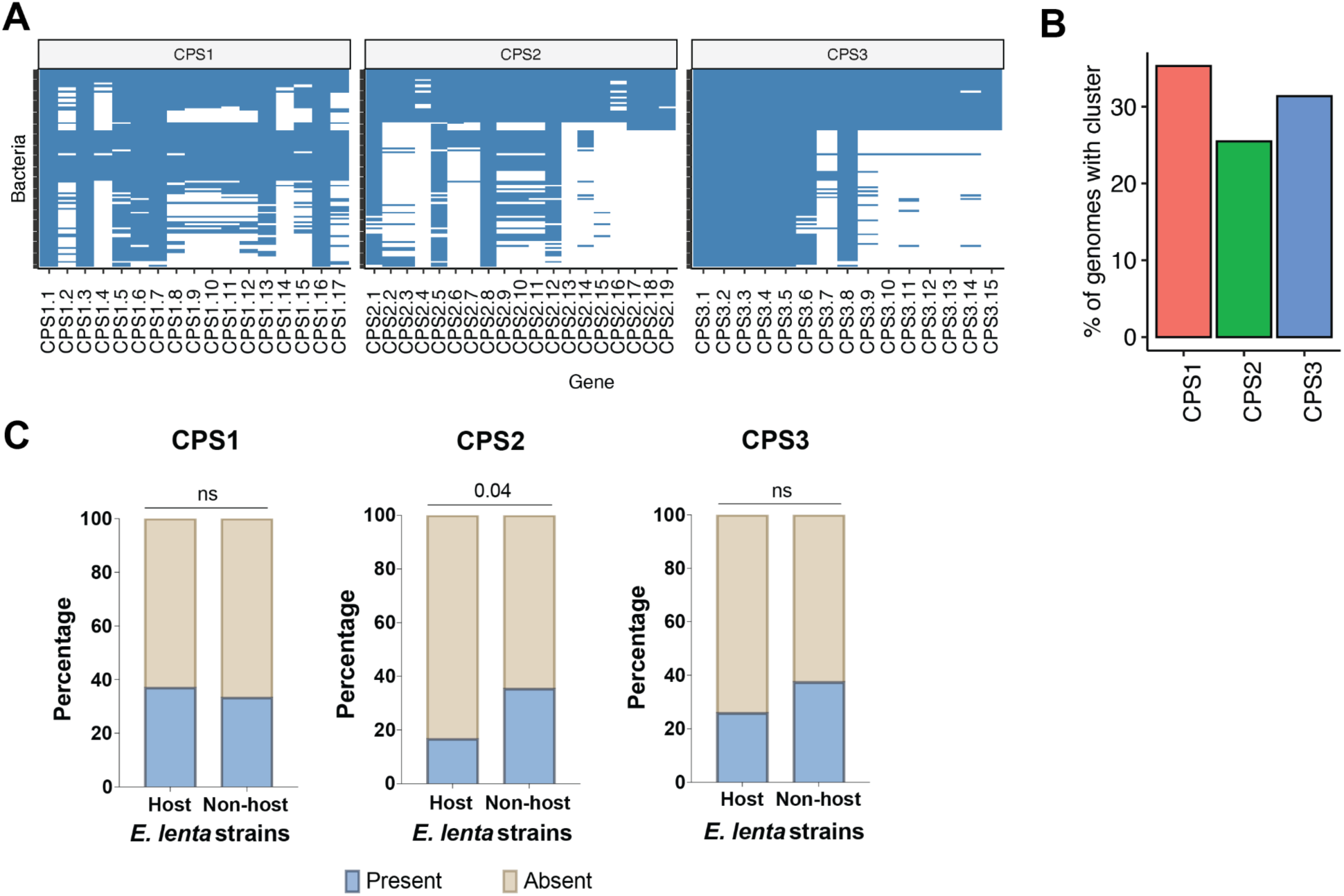
Prevalence of CPS genes in 102 *E. lenta* genomes. **(A)** Heatmap showing the proportion of genomes containing genes from each CPS cluster. **(B)** Percentage of *E. lenta* genomes harboring CPS1, CPS2, or CPS3 clusters across the strain collection. A CPS cluster was considered present if ≥85% of the genes belonging to that cluster were detected in the genome, regardless of whether the genes occurred consecutively within a single locus. **(C)** Percentage of ΦKL11 host and non-host strains containing or lacking CPS1, CPS2, or CPS3 clusters using the same ≥85% completeness criterion described in panel B. Bars represent the proportion of strains with a CPS cluster (blue) or lacking the cluster (tan). *p*-values, Fisher’s exact test; ns, not significant (*p*>0.05).

## SUPPLEMENTAL TABLES

**Table S1. Bacteriophages isolated in this study.**

**Table S2. ΦKL11 genome annotation.**

**Table S3. Gnotobiotic mouse details.**

**Table S4. *Eggerthella lenta* sequencing, assembly, and annotation summary.**

**Table S5. Structural variant (SV) details.**

**Table S6. Oligonucleotide used in this study.**

**Table S7. Genes found in each CPS cluster.**

**Table S8. Differentially expressed genes between phage-resistant and phage-sensitive *E. lenta*.**

**Table S9. Differentially expressed genes in phage-sensitive *E. lenta* in response to phage.**

**Table S10. *E. lenta* genome-wide association of orthologous group presence/absence with ΦKL11 susceptibility.**

**Table S11. *E. lenta* strains included in our comparative genomics analyses.**

## Notes

### Competing Interest Statement

The authors have declared no competing interest.

